# Recovery of consolidation after sleep following experimental stroke – interaction of slow waves, spindles and GABA

**DOI:** 10.1101/2021.01.21.427707

**Authors:** Jaekyung Kim, Ling Guo, April Hishinuma, Stefan Lemke, Dhakshin S. Ramanathan, Seok-Joon Won, Karunesh Ganguly

**Author notes:** Karunesh Ganguly, MD PhD 1700 Owens Street, San Francisco, CA, 94158, USA, Seok-Joon Won, PhD 1700 Owens Street, San Francisco, CA, 94158, USA. Mental Health Service, VA San Diego Health System, San Diego, San Diego, CA, USA. Department of Psychiatry, University of California, San Diego, San Diego, CA, USA.

## Abstract

Sleep is known to be important for promoting recovery after brain injuries such as stroke. Yet, it remains unclear how such injuries affect neural processing during sleep and how to precisely enhance sleep-dependent memory processing during recovery. Using an experimental model of focal cortical stroke in rats along with long-term electrophysiological monitoring of neural firing and sleep microarchitecture, here we show that sleep-dependent neural processing is altered after stroke induction. Specifically, we found that the precise coupling of spindles to global slow-oscillations (SO), a phenomenon that is known to be important for memory consolidation, appeared to be disrupted by a pathological increase in “isolated” local delta waves. The transition from this pathological to a more physiological sleep state – with both a reduction in isolated delta waves and increased spindle coupling to SO – was associated with sustained performance gains after task training during recovery. Interestingly, post-injury sleep processing could be pushed towards a more physiological state via a pharmacological reduction of tonic GABA. Together, our results suggest that sleep processing after cortical brain injuries may be impaired due to an increase in local delta waves and that restoration of physiological processing is important for recovery of task performance.

## INTRODUCTION

Stroke is a leading cause of long-term motor disability; despite advancements in acute management and rehabilitation, there are no widely used therapies to directly augment neural plasticity and thereby improve function (Ganguly et al., 2013; Macdonell et al., 1988; Norrving and Kissela, 2013). Importantly, it is now very clear that a major function of sleep is to regulate neuroplasticity and neural network reorganization (de Vivo et al., 2017; Genzel et al., 2014; Gulati et al., 2017; Gulati et al., 2015; Helfrich et al., 2018; Kim et al., 2019; Klinzing et al., 2019; Latchoumane et al., 2017; Ramanathan et al., 2015; Stickgold, 2005; Tononi and Cirelli, 2014; Yang et al., 2014). Thus, optimizing sleep-dependent processing during rehabilitation has the great potential to enhance recovery. While both clinical studies in human subjects and preclinical studies using various experimental models of stroke have demonstrated that sleep can influence motor recovery after stroke (Backhaus et al., 2018; Baumann et al., 2006; Duss et al., 2017; Facchin et al., 2020; Gao et al., 2010; Giubilei et al., 1992; Gottselig et al., 2002; Poryazova et al., 2015; Siengsukon and Boyd, 2009), it remains unclear precisely how sleep-dependent processing is affected by stroke.

In general, sleep-dependent processing has been closely linked to memory consolidation or the process of transforming newly encoded information into more stable long-term memories (Born et al., 2006; Stickgold, 2005). Consolidated memories are also associated with resistance to interference from learning new information (Genzel and Robertson, 2015; Robertson, 2009; Tononi and Cirelli, 2014). While consolidation is most studied in the context of hippocampal-cortical interactions (Ito et al., 2015; Molle et al., 2006; Rothschild et al., 2017; Sirota et al., 2003), sleep is also known to benefit motor memories (Kim et al., 2019; Korman et al., 2007; Lemke et al., 2021; Walker et al., 2002). Specifically, sleep-dependent processing is associated with reactivation of motor cortex neural ensembles linked to movement control and performance gains after a period of sleep (termed ‘offline’ gains) (Gulati et al., 2017; Kim et al., 2019; Ramanathan et al., 2015). Notably, memory consolidation, in general, and reactivation events, in specific, are known to require the precise coupling of sleep spindles to global slow-waves, i.e., slow-oscillations (SO) (Born et al., 2006; Buzsáki, 2015; Cairney et al., 2018; Helfrich et al., 2018; Kim et al., 2019; Latchoumane et al., 2017; Maingret et al., 2016; Ngo et al., 2013). Thus, a key goal of this study was to characterize how sleep oscillations are affected by an experimentally induced focal cortical stroke and how their characteristics may influence the recovery of sleep-associated gains in task performance, i.e., consolidation of performance.

What might be the features of sleep that are altered by an experimentally induced stroke? Recent work has shown that the balance between SO and local slow-waves – delta-waves or δ-waves – determines whether there is an effective enhancement of skill or, instead, forgetting (Kim et al., 2019). Intriguingly, neurophysiological recordings in stroke patients (Poryazova et al., 2015; Sarasso et al., 2020; Tu-Chan et al., 2017; van Dellen et al., 2013) and in animal models of stroke (Burns, 1951; Burns and Webb, 1979; Carmichael and Chesselet, 2002; Gulati et al., 2015; Nita et al., 2007) have found a prevalence of *local* low-frequency power (< 4 Hz) during awake periods; this also appears to be far more common after a cortical lesion than a purely subcortical lesion (Macdonell et al., 1988). Intriguingly, such δ-waves may also be apparent in sleep periods (Poryazova et al., 2015). Together, these observations might suggest the possibility that the sleep microarchitecture after an experimental cortical stroke contains more local δ-waves relative to global SO, thereby impairing sleep-associated “strengthening” of motor memory, i.e., bias sleep processing towards a “forget” state that is linked to weaker performance gains after sleep.

Here, we demonstrate that after induction of an experimental cortical stroke (hereafter referred to as ‘stroke_exp_’), the sleep microarchitecture of the perilesional cortex (PLC) is altered such that there is a reduced coupling of sleep spindles to SO. With the recovery of motor function with reach-to-grasp task training, there was a redistribution of sleep spindles towards a more physiological state, characterized by an increase in precise spindle-SO coupling and stronger post-training offline gains after sleep. We also found that there was a concomitant reduction in local δ-waves over this period. Interestingly, there was a direct correlation between the rate of local δ-waves and the precise coupling of spindles to SO. Remarkably, the post-stroke_exp_ imbalance between local δ-waves and global SO could be modulated by a pharmacological treatment that reduced GABA_A_-mediated tonic inhibition. Together, our results suggest that restoration of physiological sleep processing during motor recovery may allow animals to build upon each day’s training and thus allows sustained offline gains in motor performance.

## RESULTS

### Changes in spindle-SO nesting over recovery

We used an experimental stroke model to characterize how sleep processing is altered by a focal cortical lesion in rats (Clarkson et al., 2010; Ramanathan et al., 2018; Roome et al., 2014), hereafter referred to as stroke_exp_. More precisely, to examine the alterations in NREM sleep microarchitecture during recovery after stroke_exp_ (see experimental paradigm in Figure 1A), we implanted microelectrode arrays in the PLC (Ramanathan et al., 2018) (photothrombotic/PT stroke_exp_ in M1, n = 8; see animal groups in Table S1). To specifically identify phases of the motor recovery, we measured motor performance, i.e., pellet retrieval success rate in a single pellet reach-to-grasp task (Figure 1B). The reach-to-grasp task has been used as a sensitive measure of motor function; it requires reaching, grasping, and retrieving a single pellet located at a distance outside of the behavior box (Figure S1A) (Guo et al., 2021; Ramanathan et al., 2018; Whishaw et al., 1986; Wong et al., 2015). The first reach session was 1-2 weeks (9.1±2.6 days, mean±s.d.) after stroke_exp_ induction and the reestablishment of food scheduling (see Methods). Each animal experienced task training and was monitored for 6-11 sessions over a recovery period of 2-3 weeks (task performance changes over time are shown in Figure S1B for all animals). Given the variability of recovery times, each animal’s sessions were divided into tertiles; each tertile was respectively termed ‘early’, ’middle’, and ‘late’. As expected, the injury resulted in impaired task performance and there were improvements with subsequent task training. We first focused on the “pre-training” sleep (sleep period immediately prior to task training); slow-waves – SO and δ- waves – and spindles were identified using the filtered local field potential (LFP) at 0.1-4 Hz and 10-15 Hz, respectively (Figure 2A; see below for δ-waves in Figure 5; Methods). Sleep was detected using video analysis (Kim et al., 2019; Pack et al., 2007) and NREM sleep was detected using power spectral density (Figure S2; Methods). We specifically examined the changes in the temporal interactions of spindles to SO in the PLC within the pre-training sleep across sessions. The pre-training sleep likely reflects general changes in sleep microarchitecture and is less likely to reflect the immediate consequence of task training during recovery.

**Figure 1.**
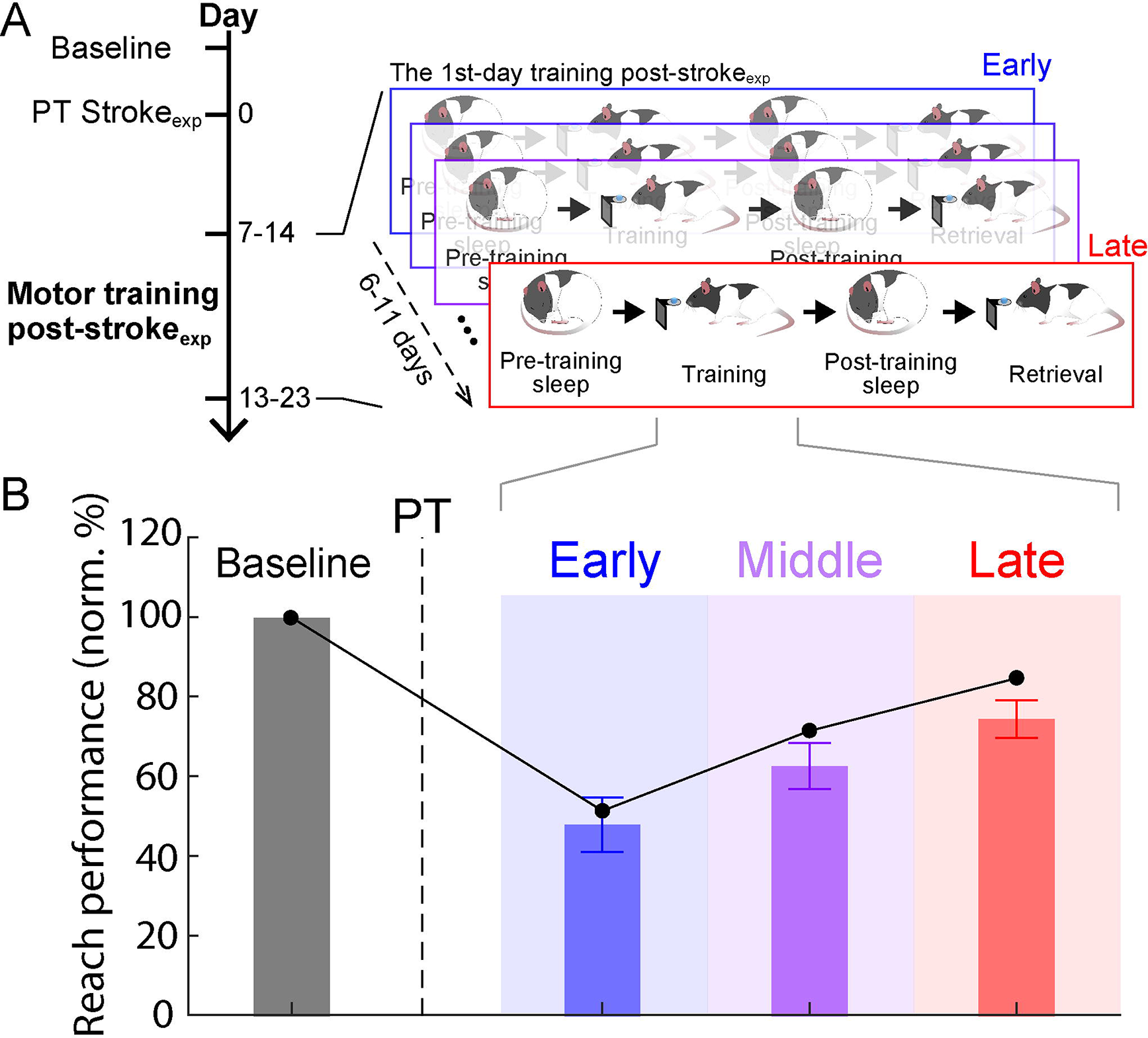
Single pellet reach to grasp task performance prior to and after an experimental stroke. **(A)**, Flow chart of motor training experiment. In a single session of the motor training post- stroke_exp_, pre-training sleep, reach training, post-training sleep, and reach retrieval blocks were monitored in sequence. PT=photothrombotic stroke_exp_. **(B)**, Mean pellet retrieval success rates (n=8 rats; black dots: two-session averages per period are shown for a single animal). Bars show average of sessions grouped into tertiles. Mean in vertical bar ± s.e.m. in errorbar. Performance was normalized by the baseline. PT stroke_exp_ impaired motor performance (baseline, n = 16 sessions in 8 rats: 100 ± 1.9% vs. early, n = 16 sessions in 8 rats: 47.8 ± 6.8%; mixed-effects model, t_30_ = –7.44, P < 10^−7^). It improved over the subsequent training (late, n = 16 sessions in 8 rats: 75.4 ± 4.7%; early vs. late, mixed-effects model, t_30_ = 5.78, P < 10^−5^).

**Figure 2.**
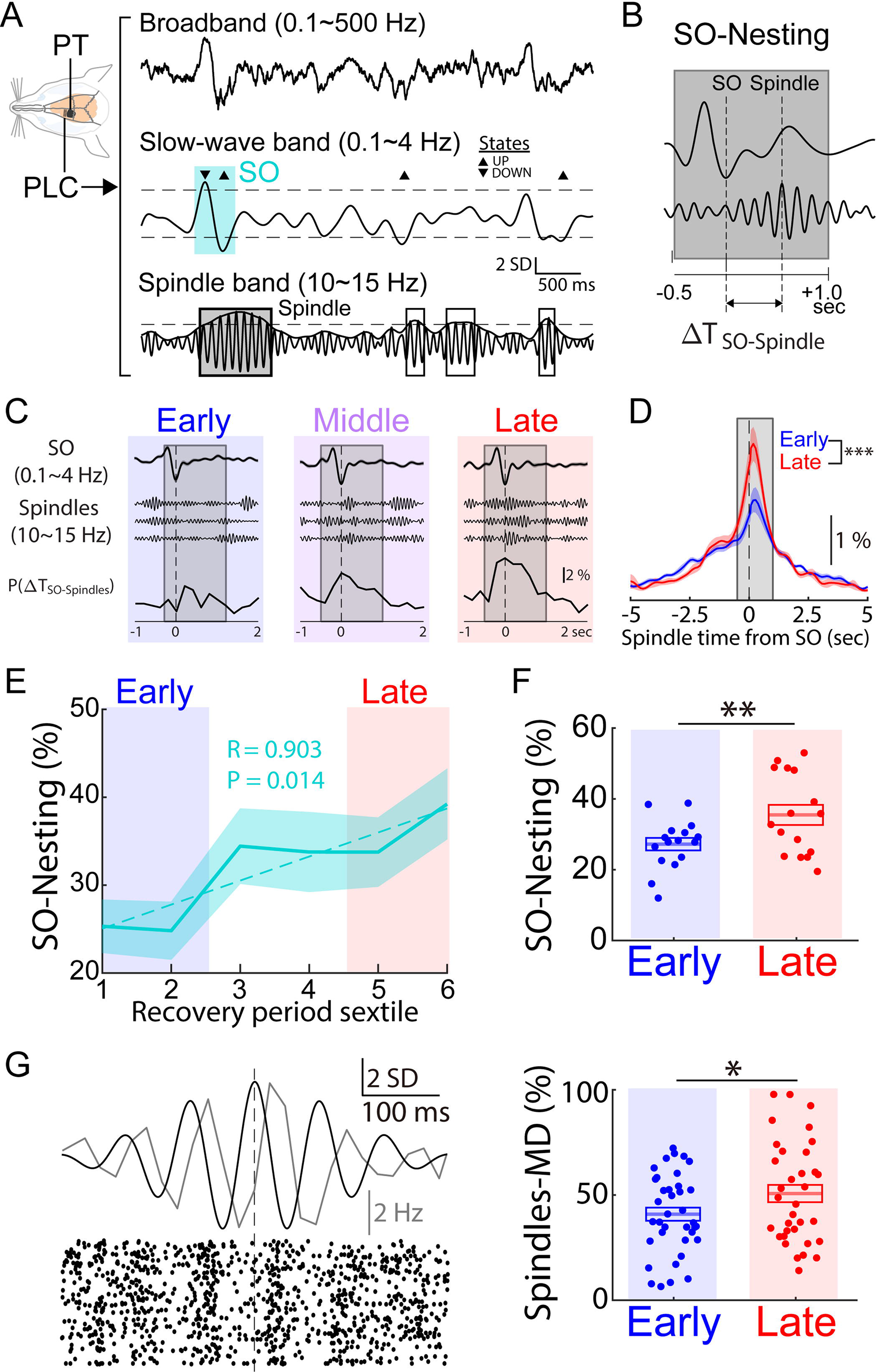
Redistribution of spindles toward SO over recovery post-stroke_exp_. **(A)**, Examples of the broadband (0.1-500 Hz) and the filtered (0.1-4 and 10-15 Hz for SO and spindles detections, respectively) trace in PLC with PT stroke_exp_. SO and spindle are marked by cyan and gray boxes, respectively. Horizental dashed lines indicate the threshold detecting SO up/down states and spindle period. Open boxes show events too brief to be labelled as a spindle. **(B)**, Cartoons of nesting of spindle to SO (SO-Nesting). Gray box indicates the time window for ‘nesting’ which is –0.5 to 1.0 sec from a SO up-state (same as ‘A’ and ‘B’). **(C)**, Examples of SO-spindle temporal interaction in each column of the respective three sessions in Figure 1B. Mean SO-up-state-triggered LFP (top, mean ± s.e.m., 0.1∼4 Hz). Three single trial examples of detected spindles near a SO (middle). Examples of a probability distribution for the spindle-peaks time from the closest SO up-state, ΔT_SO-Spindles_ (bottom). The gray window notes the nesting window. **(D)**, Comparison in distributions of ΔT_SO-Spindles_ between the early and late period (n = 16 sessions, 8 rats in each condition; Bartlett’s test, χ^2^ = 58.7, ***P < 10^−13^). **(E)**, Average time courses of the SO-Nesting and δ_I_-Nesting over the recovery period sextile (n = 8 rats). Dashed line represents linear fitting (linear regression; SO-Nesting: R = 0.903, P = 0.014). **(F)**, Comparison of SO-Nesting between early vs. late period (mean in solid line ± s.e.m. in box; SO-Nesting: early, n = 16 sessions in 8 rats, 27.2 ± 1.7% vs. late, n = 16 sessions in 8 rats, 35.5± 2.8%, mixed-effects model, t­= 3.58, **P = 0.011; using averages per period, SO-Nesting: early, n = 8 rats, 27.2 ± 2.4% vs. late, n = 8 rats, 35.5 ± 3.5%, mixed-effects model, t_14_ = 3.58, **P= 3.0 x 10^−3^). **(G)**, Left, average of event-triggered LFP (10∼15 Hz; black), spike rates (gray), and raster plot of an example unit; locked to the spindle peaks. Right, modulation-depth of units during spindles for early (n = 37 units, 8 rats; 40.9 ± 3.1%) and late period after stroke_exp_ (n = 34 units, 8 rats; 50.7± 4.1%). There were significant changes from early to late period (mixed-effects model, t_69_ = 2.21, *P = 0.030).

In the intact healthy brain, nesting of spindles to SO has been shown to be important for memory consolidation, (Antony et al., 2018; Bergmann and Born, 2018; Cairney et al., 2018; Helfrich et al., 2018; Kim et al., 2019; Latchoumane et al., 2017; Maingret et al., 2016; Ngo et al., 2013; Peyrache et al., 2011; Staresina et al., 2015). It also appears to be important for triggering offline motor performance gains (Kim et al., 2019; Lemke et al., 2021; Ramanathan et al., 2015; Silversmith et al., 2020). We thus focused on comparing the “nesting” of spindles to SO in the PLC (Figure 2B; Methods). More specifically, we measured temporal lags between a SO and its nearest spindle (ΔT_SO-Spindle_). The “SO-Nesting” focused on the events within –0.5 sec prior to and 1 sec after the peak of the up-state. With recovery, SO-Nesting became more probable (Figure 2C) and the distribution of ΔT_SO-Spindle_ became sharper, indicating spindles were occurring closer to SO (Figure 2D). Furthermore, from the distribution of ΔT_SO-Spindle_, the SO-Nesting was quantified by the probability of spindles within the nesting window (Figure 2D gray box, within a – 0.5-1s from SO up-states). SO-Nesting significantly increased over the period of motor recovery (Figure 2E). The use of a precise temporal window to define “nesting” meant that this was not simply a consequence of directly shifting spindles from one event to the other. Moreover, there was a strong increase in SO-Nesting in the late period compared to the early period (Figure 2F). We also examined the rates of SO-nested spindles and found a significant increase in the late compared to the early period (early, n = 16 sessions in 8 rats, 10.1 ± 0.60 count/min vs. late, n = 16 sessions in 8 rats, 12.3 ± 0.63 count/min, mixed-effects model, t_30_ = 2.52, P = 0.017; Figure S3). Notably, there was a significant positive relationship between the restoration of SO-Nesting and improvements in task performance (Figure 1B vs. Figure 2E; mixed-effects model fitting, R = 0.38, P = 0.031). Interestingly, SO-nested spindles were also associated with a significantly stronger spike modulation-depth/MD and amplitudes at the late compared to the early period after stroke_exp_ (MD, Figure 2G; amplitude: mixed-effects model, t_30_ = 2.26, P = 0.031, Figure S4). Taken together, these results indicate a change in the precise temporal coupling of spindles and SO during recovery.

### Changes in sleep microarchitecture directly after task training

We next wondered how task training affected the sleep microarchitecture. Past work has shown that task training in the healthy brain can induce short-term changes in spindle-SO coupling (Bergmann and Born, 2018; Cairney et al., 2018; Diekelmann and Born, 2010; Genzel et al., 2014; Helfrich et al., 2018; Kim et al., 2019; Latchoumane et al., 2017; Maingret et al., 2016; Miyamoto et al., 2017; Ngo et al., 2013; Peyrache et al., 2009; Sejnowski and Destexhe, 2000; Silversmith et al., 2020; Staresina et al., 2015), i.e., short-term *within-session* “reactive” effects of training (Figure 3A). We thus examined changes in post-training sleep “reactivity.” Interestingly, we found that the sessions within the “late” recovery period demonstrated greater increases in spindle-SO nesting following training compared to the “early” period (Figure 3B). There was not a change in the rate of SO-nested spindles (early vs. late, ΔSO-nested spindle rate: mixed-effects model, t_30_ = 1.27, P = 0.21; Figure S5). When we examined within-session changes in task performance (i.e., comparison of task performance before and after a single brief sleep period, Figures 1A and 3A), we found evidence of a significant trend in the correlation between within-session changes in SO-nesting and within-session offline gains (mixed-effect model fitting, R = 0.28, P = 0.049). Interestingly, changes in SO-Nesting were more clearly linked to changes in task performance the following day (mixed-effect model fitting, R = 0.80, P = 0.045; Figure 3C). In other words, with continued recovery time after stroke_exp_, motor training appeared to be able to trigger reactive changes in the subsequent sleep period; this appeared to manifest as improved performance the following day. This is likely consistent with recent findings that a full night of sleep results in detectable changes in functional connectivity (Lemke et al., 2021).

**Figure 3.**
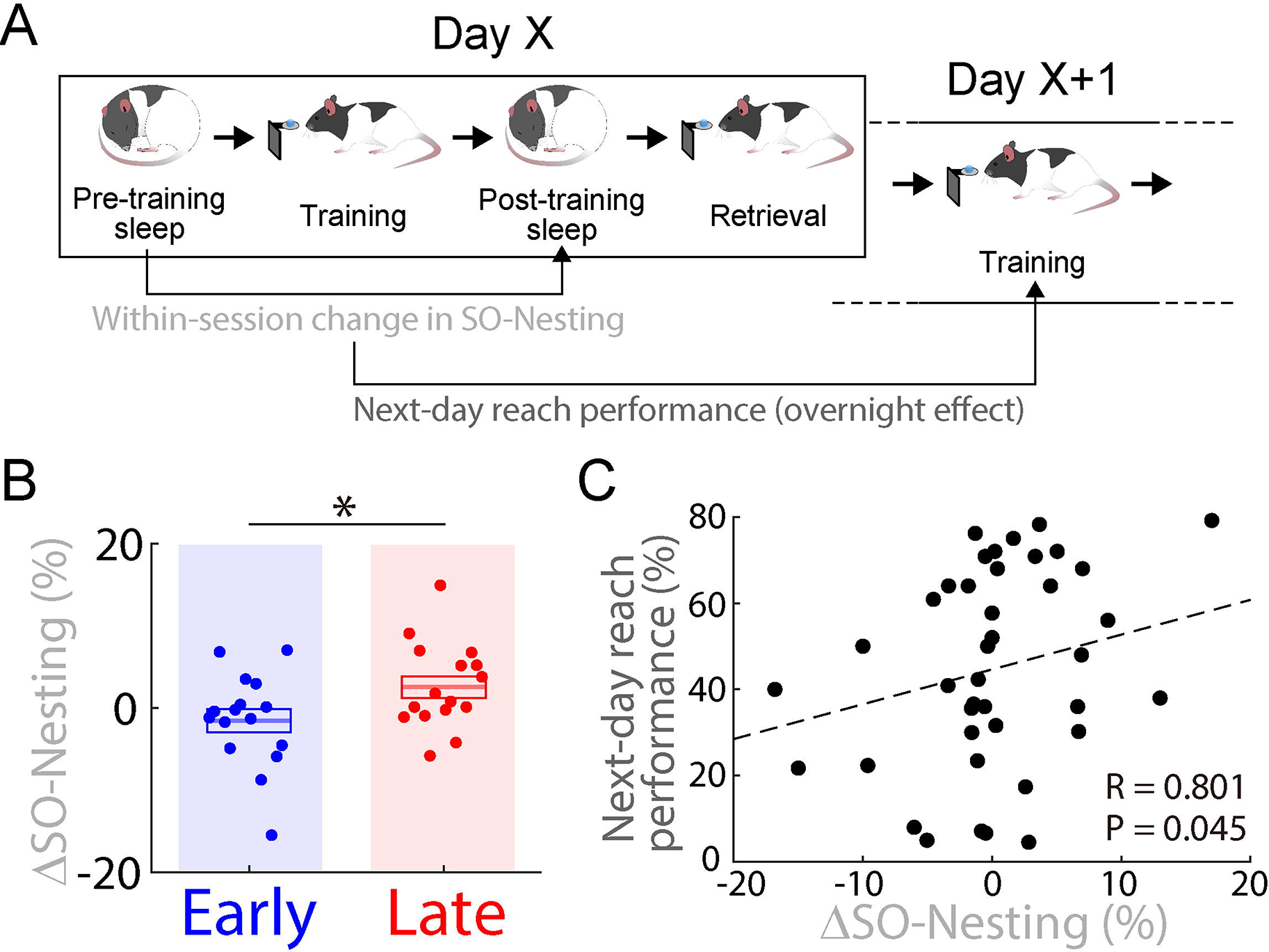
Changes in spindle redistribution and subsequent offline performance gains. **(A)**, Flow chart of experiment. Within-session changes (i.e., changes from pre-training to post- training sleep in Day X) of SO-Nesting were compared with next-day (Day X+1) reach performance. **(B)**, Comparison of changes in the SO-Nesting within a single session (ΔSO-Nesting) between the early versus the late period. ΔSO-Nesting (early, n = 16 sessions in 8 rats, –1.6 ± 1.4% vs. late, n = 16 sessions in 8 rats, 2.6 ± 1.3%, mixed-effects model, t_30_ = 2.24, *P = 0.032). Mean in solid line ± s.e.m. in box. **(C)**, Relationship of ΔSO-Nesting to the next-day reach performance over whole recovery period including early, middle, and late periods (mixed-effect model fitting, R = 0.801, P = 0.045; n = 40 sessions in 8 rats, 5 pairs in an animal).

While our previous experiments suggested that there is more robust offline gains the following day, it is also possible that there is a more direct relationship between time spent sleeping and offline gains. We thus performed behavioral experiments in separate groups with PT stroke_exp_ to test this (Figure 4). We specifically assessed the effects of sleep duration on offline motor performance gains in sessions that were in the *middle-late phase* of recovery after stroke_exp_ ; this was defined by the period in which each animal demonstrated performance improvements. In other words, we did not use *early* recovery periods when the animals showed static deficits; as shown above, this early period was linked to abnormal sleep activity. Thus, in this separate group of stroke_exp_ animals, we measured the effects of sleep on the subsequent task and compared these to a sleep restriction group (Figure 4A). For the sleep group (sleep: n = 29 sessions in 6 rats), we measured the duration of sleep using video-based detection methods. In the restriction group (no-sleep: n = 28 sessions in 8 rats), sleep was deprived either 2 or 6 hours after the training session and then the retrieval session was conducted (n = 4 restricted for 2 hours; n = 4 restricted for 6 hours). The rats with post-training sleep experienced greater offline gains, i.e., motor performance changes after post-training sleep compared to before post-training sleep, in comparison to the sleep-restricted animals (sleep: 6.4 ± 1.8%; no-sleep: –2.5 ± 2.3%; sleep vs. no-sleep, two-sample *t*-test, t_55_ = 2.28, P = 0.027). We also found a significant relationship between sleep duration and changes in task performance using the middle-late period (mixed-effect model fitting, R = 0.52, P = 0.022; Figure 4B); however, there was no significant relationship when including the early phases of recovery (mixed-effect model fitting, R = 0.25, P = 0.25). We interpreted these results to mean that sleep-associated processing might be beneficial after an initial early phase of “recovery”. Together, these results suggest a more precise dose-outcome relationship between sleep duration and the extent of post-sleep performance gains.

**Figure 4.**
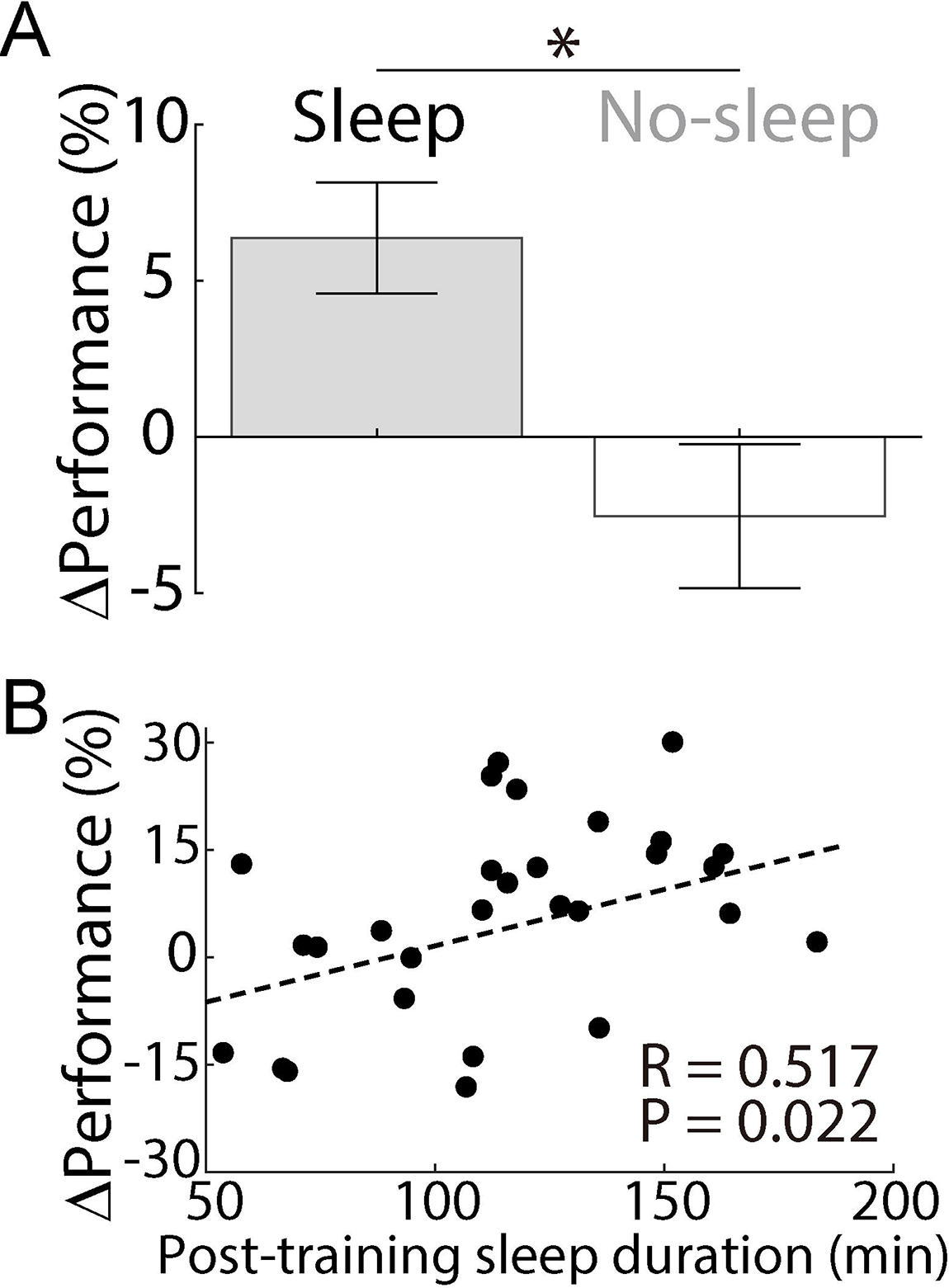
Relationship between time spent sleep post-training and motor performance. **(A)**, Within-session changes in the motor performance (ΔPerformance; changes from reach training to retrieval) in each column of sleep group and no-sleep group during the recovery after PT stroke_exp_ (mean in vertical bar ± s.e.m. in errorbar; sleep, n = 29 sessions in 6 rats: 6.4 ± 1.8%; no-sleep, n = 28 sessions in 8 rats: –2.5 ± 2.3%; sleep vs. no-sleep, mixed-effects model, t_55_ = – 2.31, *P = 0.024). In the no-sleep group, moderate vibration on the behavior box was given when rats were not active for > 40 sec, to prevent from sleeping. **(B)**, Relationship of the post-training sleep duration (general sleep by video-based detections) to the changes in motor performance for the sleep group in ‘A’ (mixed-effect model fitting, R = 0.517, P = 0.022).

### Pathologically increased local δ-waves after stroke_exp_

We next examined the changes in the temporal interactions of δ-waves with SO and spindles in the PLC. For the pre-training sleep shown in Figure 2, δ-waves were identified using the filtered LFP at 0.1-4 Hz in addition to SO and spindles (Figure 5A; Methods; SO and spindles were detected the same way in Figure 2). We then measured the temporal relationship between δ- waves and SO (ΔT_SO-δ_) in two groups of rats (PT stroke_exp_ in M1, n = 8 rats and healthy, i.e., without stroke_exp_, n = 5 rats; Figure 5B). In healthy animals, δ-waves typically follow SO; their sequential relation appears to also modulate sleep-dependent processing (Bernardi et al., 2018; Kim et al., 2019; Steriade et al., 1993; Steriade and Timofeev, 2003). We then compared it to the early period after stroke_exp_. Interestingly, in the early recovery phase, there was a significant increase in δ-waves that were apparently decoupled from SO, i.e., temporally far from the occurrence of a previous SO. All stroke_exp_ animals demonstrated this effect compared to healthy animals (Figure 5C; Figure S6 compares the respective distributions of ΔT_SO-δ_ before and after stroke_exp_ in the same animal). Using this distribution, we separated δ-waves into two classes, i.e., “SO-coupled/δ_SO_” and “isolated/δ_I_”. This distinction was based on simply dividing the ΔT_SO-δ_ distributions using the 50^th^ percentile; δ-waves less than the 50^th^ percentile were labeled δ_SO_, whereas the remaining δ-waves were labeled δ_I_ (Methods). Notably, the overall ratios of SO/δ were similar in both stroke_exp_ and healthy animals (stroke_exp_: 27.9 ± 2.2%; healthy: 32.4 ± 2.6%; stroke_exp_ vs. healthy, two-sample *t*-test, t_23_ = –1.31, P = 0.24). Thus, after stroke_exp_, while the relative number of SO and δ-waves were comparable to the healthy state, there was, however, a significant increase in the prevalence of local δ_I_-waves that were not temporally linked to global SO. We also examined the changes in the temporal interactions of spindles δ_I_-waves in the PLC. We analyzed “δ_I_-Nesting” in a similar manner to “SO-Nesting” illustrated in Figure 2, i.e., for δ_I_- waves and its closest spindle (Figure 5D; Methods). With recovery, δ_I_-Nesting significantly decreased over the period of motor recovery (Figure 5E). Moreover, there was a moderate decrease in the late period compared to the early period (Figure 5F). We also examined the rates of δ_I_-nested spindles and found no significant change (early, n = 16 sessions in 8 rats, 10.3 ± 0.70 count/min vs. late, n = 16 sessions in 8 rats, 11.1 ± 0.51 count/min, mixed-effects model, t_30_ = 1.18, P = 0.25; Figure S3). Notably, there was a negative relationship between the decrease of δ_I_-Nesting and time-dependent improvements in task performance (Figure 1B vs. Figure 5E; mixed-effects model fitting, R = –0.43, P < 10^−2^); this was opposite of the relationship we found above for SO-Nesting. The relative nesting, i.e., SO-Nesting/δ_I_-Nesting was more predictive of the recovery of task performance compared to the SO-Nesting (SO-Nesting/δ_I_-Nesting: mixed-effects model fitting, R = 0.46, P < 10^−2^; smaller absolute errors between the true and the predicted performances by SO-Nesting/δ_I_-Nesting versus SO-Nesting: paired *t*-test, t_31_ = 2.18, P = 0.037). Taken together, these results indicate there was a change in the precise temporal coupling of spindles toward SO and away from δ_I_-waves with recovery.

**Figure 5.**
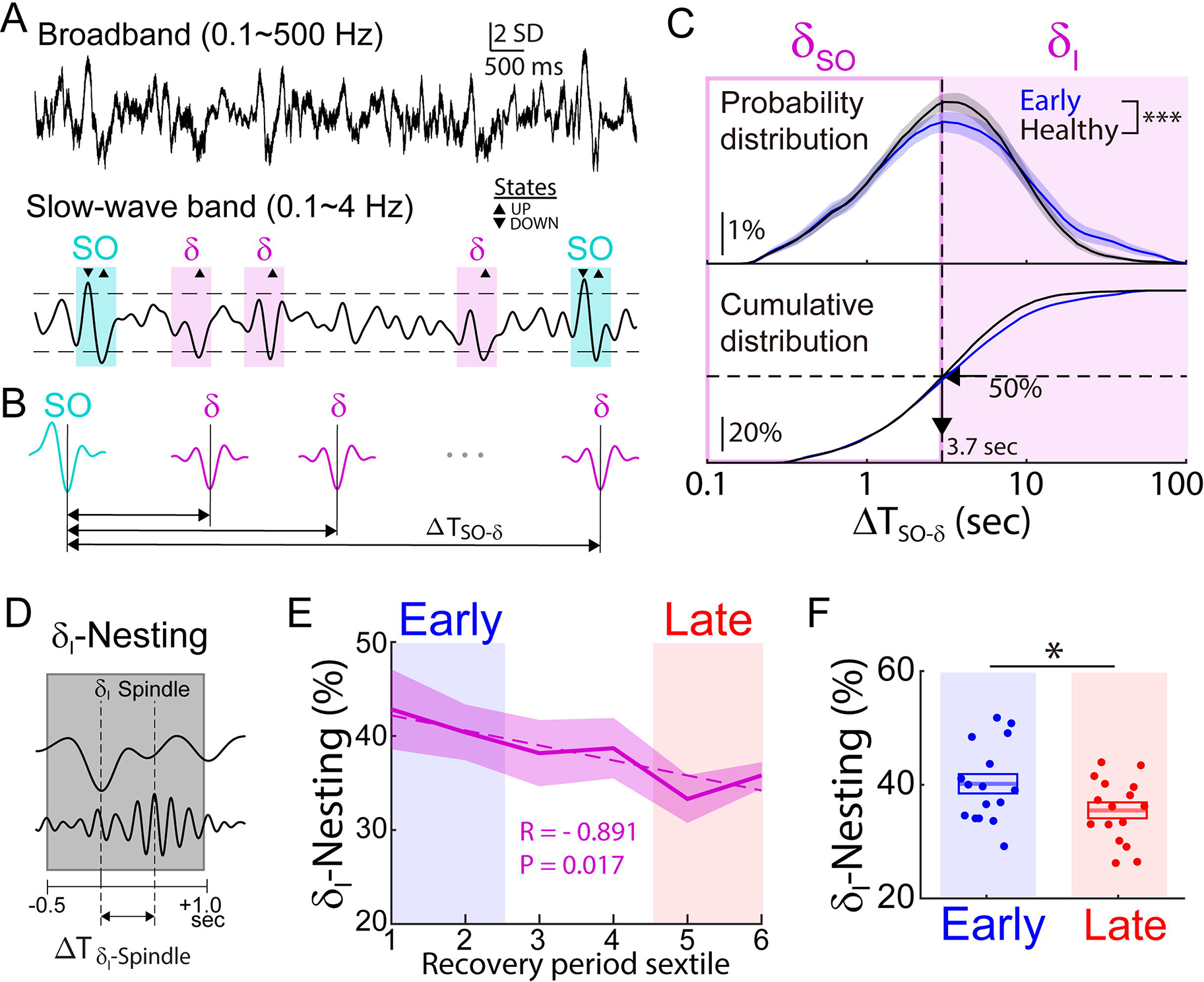
Pathological increase of isolated δ (δ_I_) after stroke_exp_. **(A)**, Examples of the broadband (0.1-500 Hz) and the filtered (0.1-4 Hz) trace showing SO and δ- waves in PLC. SO and δ-waves are marked by cyan and magenta boxes, respectively. **(B)**, Schematic showing the temporal distance of δ-waves from SO (ΔT_SO-δ_). **(C)**, Comparison of SO-coupled δ (δ_SO_) and isolated δ (δ_I_) waves for sleep sessions in the early period (blue; n = 16 sessions, 8 rats) and sleep sessions in the intact brain of healthy animals. (black; n = 9 sessions, 5 rats; Kolmogorov-Smirnov test, KS-statistic = 0.17, ***P < 10^−15^). Average distributions of ΔT_SO-δ_ are shown. δ_SO_ and δ_I_ waves were separated by the 50th percentile (3.7 sec, vertical dashed line) from the distribution with healthy animals. **(D)**, Cartoon of nesting of spindle to δ_I_ (δ_I_-Nesting). Gray box indicates the time window for ‘nesting’ which is –0.5 to 1.0 sec from a δ_I_ up-state. **(E)**, Average time courses of the δ_I_-Nesting over the recovery period sextile (n = 8 rats). Dashed line represents linear fitting (linear regression; δ_I_-Nesting: R = –0.891, P = 0.017). **(F)**, Comparison of δ_I_-Nesting between early vs. late period (mean in solid line ± s.e.m. in box; δ_I_- Nesting: early, n = 16 sessions in 8 rats, 40.1 ± 1.7% vs. late, n = 16 sessions in 8 rats, 35.4 ± 1.4%, mixed-effects model, t_30_ = –2.27, *P = 0.031; using average per period, δ_I_-Nesting: early, n = 8 rats, 40.1 ± 2.3% vs. late, n = 8 rats, 35.4 ± 1.7%, mixed-effects model, t_14_ = –1.74, **P = 0.10).

### Relationship between local δI-waves preceding SO and nesting

Is there a more direct relationship of these observed changes in δ_I_-waves on SO-spindle nesting? Past studies have shown that changes in SO-associated parameters (e.g., amplitude and refractoriness in spindle generating networks) can depend on the temporal proximity of slow-waves (Bernardi et al., 2018; Ngo et al., 2015). Given the changes we observed in SO-Nesting with recovery, we thus examined whether changes in how often δ_I_-waves directly preceded a SO event could predict this change. We specifically examined how many δ_I_-waves preceded SO, i.e., –5-0s from a SO up-state (Figure 6A). Remarkably, this phenomenon appeared to significantly influence spindle nesting to SO; we found a strong negative correlation between the δ_I_-wave rate preceding SO and SO-Nesting (Figure 6B). Furthermore, the rate of δ_I_-waves preceding SO had a significant negative correlation with the task-performance the following day (Figure 6C) as well as the within-session changes in task performance (mixed-effect model fitting, R = 0.30, P = 0.032). Thus, our results suggest that the prevalence of δ_I_-waves in the early phase may influence the nesting of spindles to SO; the prevalence of δ_I_-waves was also correlated with changes in task performance.

**Figure 6.**
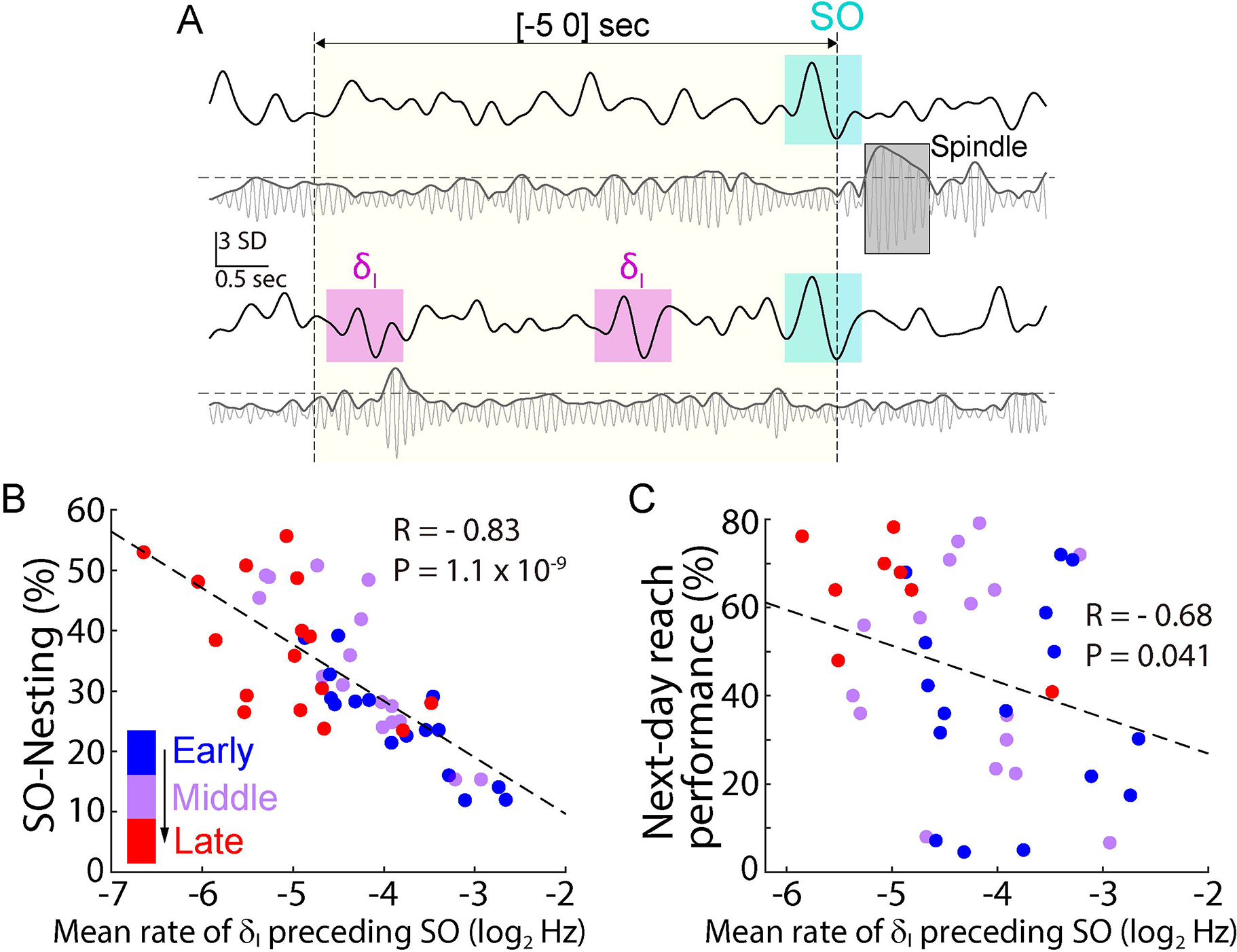
Effect of δ_I_ preceding SO on SO-spindle nesting. **(A)**, Top, example LFP traces filtered at slow-wave band (0.1-4 Hz; top) and spindle band (10-15 Hz; bottom) in PLC for a case where no δ_I_-wave preceded SO during –5-0s from a SO up-state (yellow box). SO, δ_I_-waves, and Spindle were detected the same way shown in Figures 2 and 5. Horizontal dashed line indicates the threshold for detecting spindles. Bottom, same as ‘top,’ but for a case where two δ_I_-waves preceded SO during –5-0s from a SO up-state. **(B)**, Relationship between the mean rate of δ_I_-waves preceding SO within 5 sec and the SO-Nesting over all recovery sextiles. There was a negative correlation (n = 48 sessions in 8 rats, 6 pairs in an animal; dashed line, mixed-effect model fitting, R = –0.83, P = 1.1 x 10^−9^). This suggests that the prevalence of δ_I_-waves directly preceding SO strongly modulates SO-spindle nesting. **(C)**, Relationship between the mean rate of δ_I_-waves preceding SO within 5 sec to the next-day reach performance in Figure 3C over all recovery sextiles. There was a negative correlation (n = 40 sessions in 8 rats, 5 pairs in an animal; dashed line, mixed-effect model fitting, R = –0.68, P = 0.041).

### Reducing elevated tonic inhibition and pathological sleep

What might be the underlying neurophysiological basis for our observed changes in SO and δ_I_- waves with recovery? While both spontaneous changes in structural connectivity over time and motor training might contribute, another intriguing possibility is that the elevations in extra-synaptic γ-aminobutyric acid (GABA) after experimental stroke may alter local excitability and thus perturb the ability of global SO to organize local δ-waves. Studies have reported that elevated GABA_A_- mediated tonic inhibition reduces network excitability, and that reducing such inhibition can have a beneficial effect on recovery after stroke_exp_ (Clarkson et al., 2010; He et al., 2019). These studies specifically imply that an altered excitation-inhibition/E-I balance (Kim and Fiorillo, 2017; Xia et al., 2017; Yizhar et al., 2011) plays an important role in motor recovery.

Thus, we tested the effects of blocking GABA_A_-mediated tonic inhibition after stroke_exp_ on the temporal coupling of spindles to slow-waves. For these experiments, we only tested this intervention during the early spontaneous recovery period; this allows us to examine changes in the absence of contributions from task training. We specifically assessed the effects of injecting the GABA_A_ α5-subtype receptor inverse agonist (L655-708) (Clarkson et al., 2010) during early recovery after a PT stroke_exp_ (n = 3 rats) and an Endothelin-1(ET-1) induced stroke_exp_ (n = 3 rats); for the ET-1 induced stroke_exp_ microelectrode arrays were targeted to the PLC; an attached cannula was used for infusion of ET-1 into M1 (Methods). The ET-1 approach allowed us to collect baseline neural data prior to stroke induction; pre-stroke_exp_ baseline spiking/LFP activity was recorded only in the ET-1 induced stroke_exp_ animals. This was then followed by recordings after the injection of vehicle (sham) or L655-708 (drug) during the early phase of recovery, i.e., days 7∼12 after stroke_exp_ in both stroke_exp_ models (Figure 7A; Methods).

**Figure 7.**
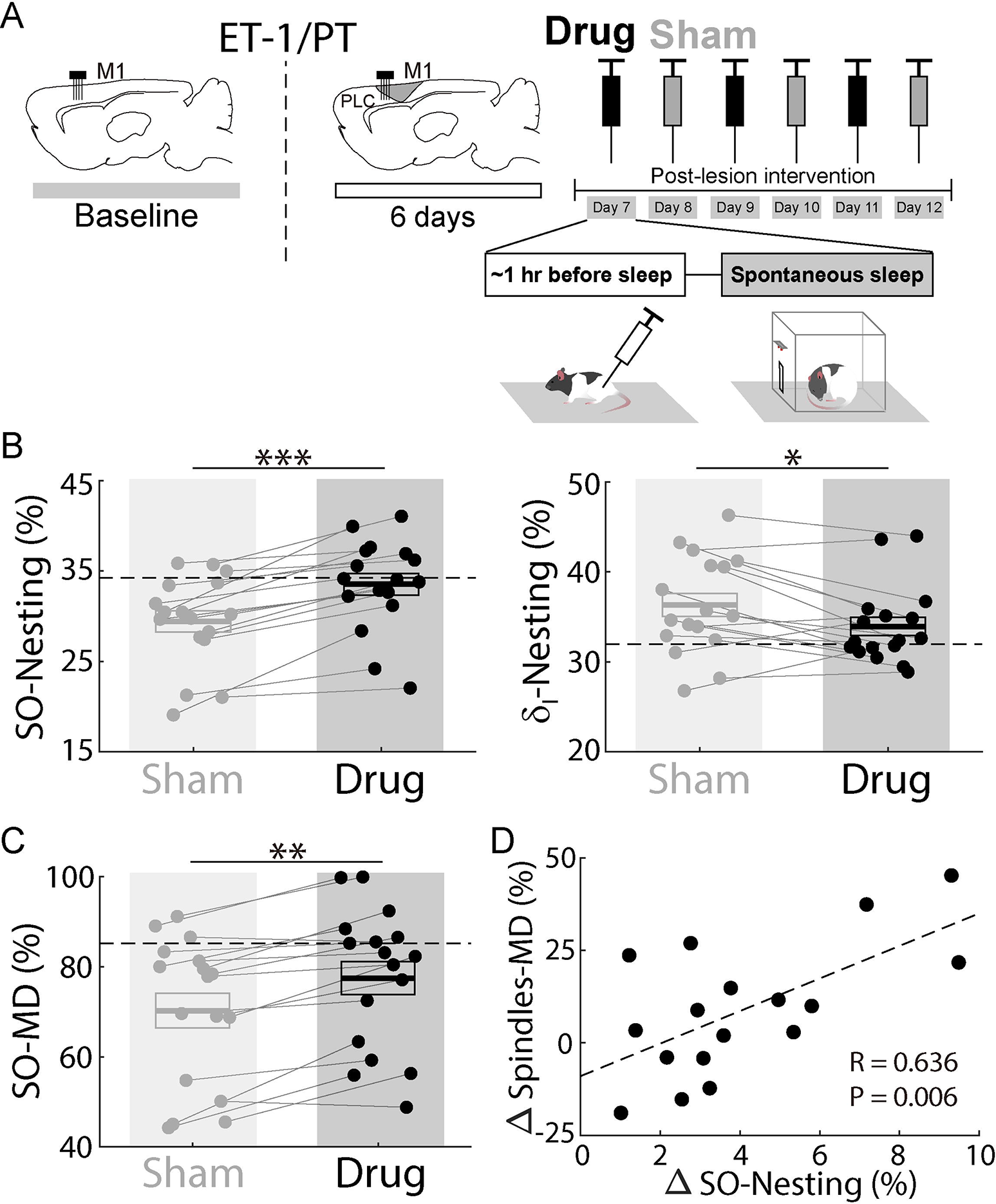
Blocking GABA_A_ α5-subtype receptor increases SO-Nesting. **(A)**, Flow chart of experiment blocking tonic GABA_A_. 32-channel microelectrode arrays were implanted into PLC to monitor LFP/spike (n = 6 rats; three rats for ET-1 induced and three rats for photothrombotic model in M1). In the only ET-1 induced stroke_exp_, LFP/spike activity was recorded 2∼3 days before inducing stroke_exp_ (vertical dashed line) for the baseline (horizontal dashed lines in “B” and “C.” Either GABA_A_ α5-subtype receptor inverse agonist (L655-708; drug) or vehicle (saline; sham) was administrated IP on separate days, i.e., the drug condition was tested every other day during range of day 7-12. The starting condition was the drug at day 7 in three rats and the sham at day 7 in the other three rats. **(B)**, Comparison of SO-Nesting and δ_I_-Nesting between the drung and the sham conditions. SO-Nesting and δ_I_-Nesting was stronger and weaker, respectively, with the drug compared to the sham (SO-Nesting: sham, n = 17 sessions, 29.4 ± 1.2% vs. drug, n = 17 sessions, 33.5 ± 1.2%, mixed-effects model, t_32_ = 6.93, ***P < 10^−7^; δ_I_-Nesting: sham, 36.3 ± 1.3% vs. drug, 33.9 ± 1.0%, mixed-effects model, t_32_ = –2.29, *P = 0.028). Baseline in horizontal dashed line, mean in solid line ± s.e.m. in box (also in “C”). **(C)**, Comparison of unit modulation-depth during SO, i.e., SO-MD; (maximum - minimum) firing rates within –0.5 to 0.3 sec over baseline firing rates within –2 to –0.5 sec from SO up-states. SO-MD was higher with the drug compared to the sham (sham, 70.3 ± 3.8% vs. drug, 77.5 ± 3.7%, mixed-effects model, t_32_ = 2.79, **P < 0.01). **(D)**, Relationship from the changes of SO-Nesting in ‘B’ to the changes of modulation-depth during spindles, i.e., spindles-MD; calculated like SO-MD in ‘C’ but within –0.25 to 0.15 sec from spindles peaks. There was a positive significant correlation (mixed-effect model fitting, R = 0.636, P = 0.006).

Strikingly, treatment with the drug for these stroke_exp_ animals resulted in significantly stronger SO-spindle nesting with a concomitant reduction in δ_I_-spindle nesting (Figure 7B). Notably, drug infusion in a separate group of healthy animals resulted in stronger δ_I_-spindle nesting with non-significant changes in SO-spindle nesting (Figure S7; SO-Nesting: sham, n = 9 sessions in 3 rats, 28.4 ± 0.73% vs. drug, n = 9 sessions in 3 rats, 26.8 ± 1.7%, mixed-effects model, t_16_ = –1.14, P = 0.27; δ_I_-Nesting: sham, 35.2 ± 1.1% vs. drug, 38.7 ± 1.3%, mixed-effects model, t_16_ = 3.59, P < 10^−2^). It suggests that the beneficial effect of the drug is specific for stroke_exp_ animals while it is detrimental to physiological processing in healthy animals (see also Discussion). In stroke_exp_ animals, we also observed increased excitability when comparing the drug versus sham. For example, the modulation-depth of cortical spiking during SO was stronger with the drug (Figure 7C). Moreover, changes in the modulation-depth of cortical spiking during spindles, a measure of the effectiveness of spindles in modulating local spiking, could be explained by an increase in SO-nesting (Figure 7D). We did not find any differences in effects for the ET-1 and PT groups (ET-1 vs. PT; ΔSO-Nesting: two-sample *t*-test, t_15_ = –0.14, P = 0.89; ΔSpindles-MD: two-sample *t*-test, t_15_ = –0.064, P = 0.95). As noted above, the baseline level was measured in the ET-1 group, and also compared with the two experimental conditions. For the SO-spindle nesting and δ_I_- spindle nesting, the sham showed significant differences compared to the baseline, i.e., drop in SO-spindle nesting and increase in δ_I_-spindle nesting after stroke_exp_, while the drug was not significantly different from the baseline (SO-Nesting: sham vs. baseline, mixed-effects model, t_13_ = 3.93, P = 1.7 x 10^−3^, drug vs. baseline, mixed-effects model, t_13_ = 2.01, P = 0.067; δ_I_-Nesting: sham vs. baseline, mixed-effects model, t_13_ = –3.09, P = 8.6 x 10^−3^, drug vs. baseline, mixed-effects model, t_13_ = –1.55, P = 0.15). Thus, our results indicate that elevated GABA_A_-mediated tonic inhibition can alter the sleep microarchitecture towards an apparently more physiological state in the early recovery period.

## DISCUSSION

Overall, our results reveal how sleep-associated processing is affected by a focal cortical stroke_exp_ and how such processing interacts with task training during motor recovery (Figure 8). First, our results indicate that, after induction of stroke_exp_, there is an alteration in the precise coupling of spindles to SO and that such changes are correlated with overnight offline gains in task performance during recovery. Second, we found that after stroke_exp_, the sleep microarchitecture is characterized by an increase of local δ_I_-waves relative to the healthy intact brain. Such increases in local δ_I_-waves were also linked to changes in SO-spindle coupling and offline gains. Third, our results suggest that pharmacological modulation of extracellular tonic GABA after stroke_exp_ could both significantly improve the precision of SO-spindle coupling as well as reduce local δ_I_-waves. Together, these suggest that known changes in E-I balance after stroke_exp_ could impair normal sleep processing and also reduce the extent of offline performance gains following task training during recovery. More broadly, our results suggest a roadmap to delineate normal vs pathological sleep after stroke; they also suggest novel therapeutic targets to modulate sleep-associated processing (e.g., tonic GABA and SO-spindle nesting) in order to enhance motor recovery.

**Figure 8.**
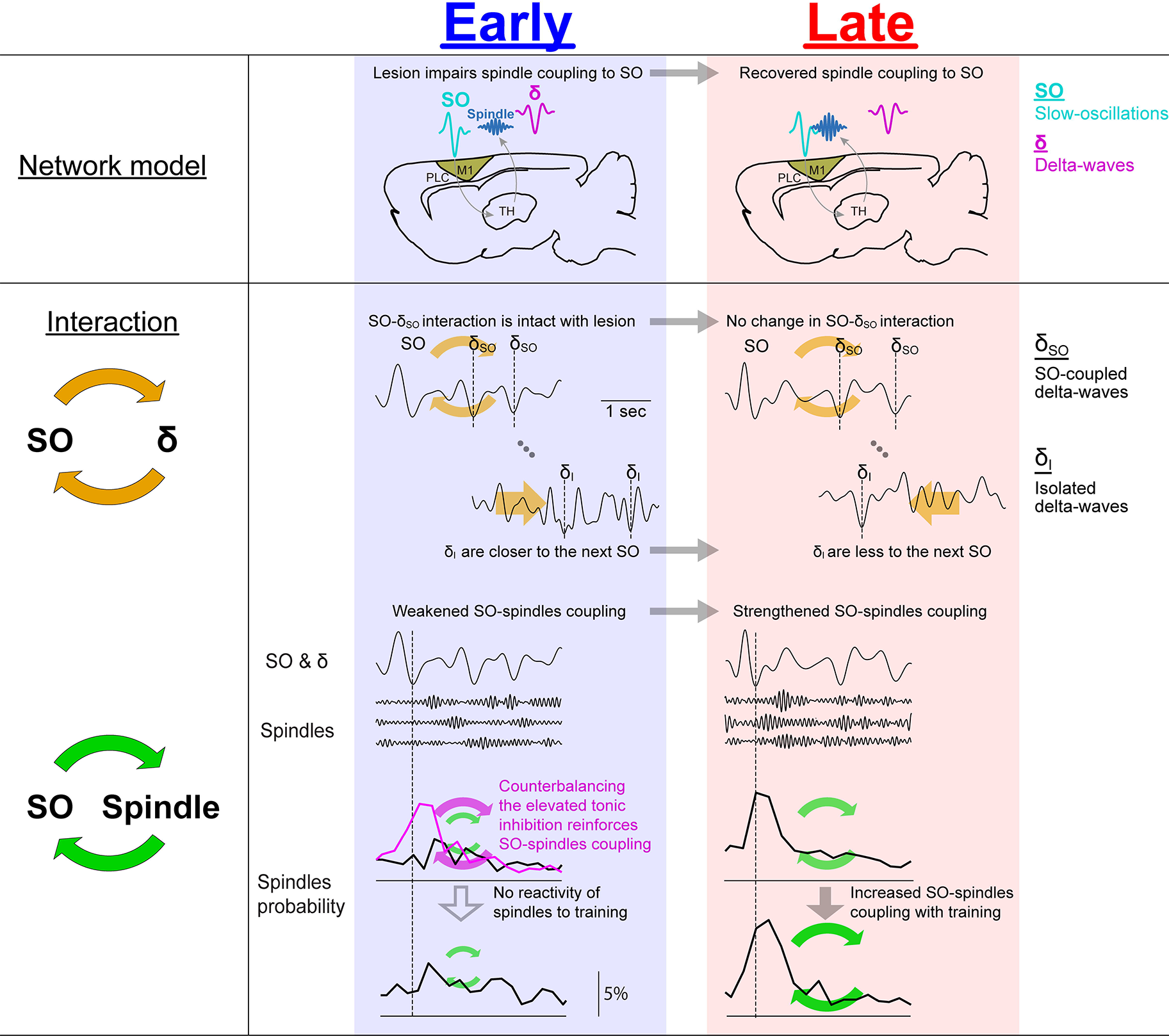
Schematic summary for the proposed model and main results. Stroke_exp_ in brain likely impairs thalamocortical spindle coupling to SO (blue box, early) and it is more normal with recovery, i.e., precise locking to SO at the late recovery period (red box). Interactions between SO and SO-coupled δ-waves (δ_SO_) is intact after stroke_exp_, while δ-waves that are isolated from SO (δ_I_) are pathologically redistributed away from SO; pathological δ_I_-waves reduces with recovery. SO-spindle coupling is weakened with stroke_exp_ (early period) and it is normalized with long-term training-induced recovery (across-session). Spindles are not reactive to the motor training (within-session) in early period, i.e., no change in SO-spindle coupling from pre-training sleep to post-training sleep. However, SO-spindle nesting is more reactive to the training with continued recovery. Remarkably, pharmacological treatment to counterbalance the presumed elevated tonic inhibition after stroke_exp_ may drive memory consolidation and, therefore, enhance SO-spindle coupling associated with functional motor recovery after stroke_exp_.

### Link between sleep microarchitecture and memory consolidation post-stroke

EEG studies in stroke patients (Macdonell et al., 1988; Poryazova et al., 2015; Tu-Chan et al., 2017; van Dellen et al., 2013) and LFP recordings in animal models of stroke_exp_ (Carmichael and Chesselet, 2002; Gulati et al., 2015) have found evidence of increased low-frequency power during awake periods after stroke. These studies have also indicated that this increased awake low-frequency power is a marker of cortical injury and loss of subcortical inputs post-stroke (Topolnik et al., 2003). Our findings of increased local δ_I_-waves during post-stroke_exp_ sleep are perhaps related to this phenomenon. Importantly, the increased frequency of δ_I_-waves preceding SO was closely associated with the attenuated temporal coupling of spindles to SO (Figure 6). We further found evidence for a normalization of SO-δ_I_ interactions towards a physiological state with recovery and task training.

In general, there is growing evidence that the temporal coupling of spindles to SO are essential drivers of memory consolidation during sleep (Antony et al., 2018; Cairney et al., 2018; Helfrich et al., 2018; Kim et al., 2019; Latchoumane et al., 2017; Maingret et al., 2016; Staresina et al., 2015). Such coupling has been linked to spike-time dependent plasticity (Bergmann and Born, 2018). It is also a potent driver of the reactivation of awake experiences (Antony et al., 2018; Cairney et al., 2018; Ngo et al., 2013; Peyrache et al., 2009). Finally, disruption of this coupling has been found to impair memory consolidation after awake experiences (Kim et al., 2019). Thus, our results suggest a link between pathological changes in network physiology after stroke_exp_ to impaired sleep-associated processing, primarily via disruption of precise spindle-SO coupling.

It is important to note that our observations of changes in sleep microstructure during recovery are largely correlational at this stage. It is difficult for us to distinguish the effect of a deficient capacity of encoding during awake training from our observed changes in sleep processing and offline gains following sleep. For example, it is likely that with recovery, more effective task performance and associated ensemble dynamics (Guo et al., 2021; Khanna et al., 2021; Ramanathan et al., 2018) also influence the efficacy of sleep processing. However, it is worth pointing out that precise manipulations of sleep processing in healthy intact animals are sufficient to prevent offline gains even when awake task learning/performance was robust. For example, our recent study found that relatively precise modulation of the extent of sleep spindle-SO coupling in healthy animals could either enhance or impede sleep processing (Kim et al., 2019). Future studies using such precise modulation approaches can causally test whether manipulation of sleep processing after stroke_exp_ is sufficient to enhance or impede motor recovery. Such an approach can also more precisely determine how awake task training during recovery influences sleep processing.

### Implications for rehabilitation

We found that changes in sleep-associated processing after stroke_exp_ were correlated with offline task performance gains duringrecovery. Importantly, clinical rehabilitation approaches are quite varied and not simply equal to task training; moreover, there are differences in training that aim to restore function as opposed to enabling functional compensation (Bernhardt et al., 2017; Ganguly et al., 2013; Pearson-Fuhrhop et al., 2009). Even so, the reach training used in this study does offer a way to study functional recovery after stroke_exp_ in both rodents and non-human primates (Guo et al., 2021; Khanna et al., 2021; Ramanathan et al., 2018; Whishaw et al., 1986). Based on this, our results specifically suggest that marking whether sleep is in a pathological (i.e., local δ_I_-waves increase and weak SO-spindle coupling) or in a physiological state may be an important consideration for the timing of intense rehabilitation. Our current results, however, do not allow us to determine whether this transition is specifically an early benefit of attempted motor training or a process that is independent of training. Interestingly, past studies in humans and rodents have suggested a sensitive period in which training can trigger long-term benefits (Dromerick et al., 2021; Dromerick et al., 2009; Krakauer et al., 2012). Notably, if it is indeed the case that awake low-frequency power in stroke patients is related to our observed effects on sleep processing, it might help explain past results indicating that awake low-frequency power is a predictor of recovery. Future studies using EEG can measure in greater detail how changes in the coupling of spindles to SO and δ_I_-waves predict the benefits of task training. Such studies can also help delineate when and if the transition to physiological sleep occurs and whether this is at all related to the sensitive period; this is particularly important as the time course of recovery is much longer in humans than in rodents.

### Implications for therapeutic neuromodulation

Our results have implications for future translational studies. Animal studies have suggested that GABA_A_-mediated tonic inhibition in the PLC may be a therapeutic target to promote recovery in the acute/early phases after stroke; blockade of GABA_A_-mediated tonic inhibition was found to promote motor recovery maximally within the first 1∼2 weeks in mice (Clarkson et al., 2010; He et al., 2019). Both short-term (i.e., acutely prior to task training) and long-term (i.e., chronic infusion) of the drug have been tested. Intriguingly, long-term infusion appears to be much better (Clarkson et al., 2010). Although our pharmacological experiments were performed in a separate group of animals without motor training, our results provide a possible mechanism for why long-term infusion by Clarkson and colleagues may be essential to achieve prominent benefits; more specifically, the effect of this drug might not only help with task-specific online training, i.e., encoding motor skills during awake training, but also promote offline memory consolidation during sleep. Future work can better define the causal role of drug manipulation during sleep in the recovery of motor function after stroke. An important note for the therapeutic approach is the timing of drug delivery. A past study using a mouse stroke model pointed out that excessive increases in excitability via modulation of GABA_A_-mediated tonic inhibition could exacerbate injury after stroke (Clarkson et al., 2010). Interestingly, our test of this drug in healthy animals resulted in the opposite modulation of spindle nesting; spindles were coupled more strongly with δ_I_-waves compared to the sham condition (Figure S7). Together, it suggests that the timing of neuromodulation is important. Such knowledge is likely to be critical for successful translation to clinical populations.

Our results also suggest possible approaches to neuromodulation to enhance recovery. It is quite likely that SO and local δ-waves can be monitored using non-invasive EEG recording in clinical stroke patients, ideally allowing patient-specific therapeutics. For example, non-invasive brain stimulation during sleep on memory consolidation (Antony et al., 2018; Cairney et al., 2018; Marshall et al., 2006; Ngo et al., 2013) can be tailored based on suppressing δ-waves waves or enhancing SO. It is also possible that invasive approaches for neuromodulation after stroke (Levy et al., 2016) can also focus on optimizing sleep-associated processing. For example, while recent studies have shown that direct electrical stimulation can enhance awake performance after stroke_exp_ (Khanna et al., 2021; Ramanathan et al., 2018), there has been less of a focus on extending this to sleep periods. Interestingly, a recent study suggested that neuromodulation to modulate up-states during sleep can enhance recovery after stroke_exp_ (Facchin et al., 2020). It is quite possible that closed-loop neuromodulation approaches that focus on both optimizing task performance and its consequence during sleep-associated processing (Kim et al., 2019) may lead to the greatest long-term benefits during rehabilitation after stroke.

## METHODS

### Animals/surgery

Experiments were approved by the Institutional Animal Care and Use Committee at the San Francisco VA Medical Center. We used a total of twenty eight adult Long-Evans male rats (250– 400 g; Charles River Laboratories); twenty six rats with focal experimental stroke (stroke_exp_) and two rats without stroke_exp_ (Table S1; detailed contributions of each rat on the respective results). No statistical methods were used to predetermine sample sizes, but our sample sizes are similar to those reported in previous publications (Gulati et al., 2017; Gulati et al., 2014; Ramanathan et al., 2015). Animals were kept under controlled temperature and a cycle with 12-h light and 12-h dark (lights on at 06:00 a.m.). Surgeries were performed under isofluorane (1-3%) anesthesia and body temperature was maintained at 37 °C with a heating pad. Atropine sulfate was also administered intraperitoneal/IP before anesthesia (0.02 mg/kg of body weight). We implanted 32/128-channel microwire arrays for recording LFP/spike activity; arrays were lowered down to 1,400-1,800 μm in layer 5 of the primary motor cortex (M1) or perilesional cortex (PLC) in the upper limb area. In the healthy animals (n = 2), neural probes were centered over the forelimb area of M1, at 3 mm lateral and 0.5 mm anterior from the bregma. In the stroke_exp_ animals (n = 14), the neural probe was placed immediately anterior to the lesion site, typically centered ∼3-4 mm anterior and 2.5-3 mm lateral to the bregma. The reference wire was wrapped around a screw inserted in the midline over the cerebellum. The final localization of depth was based on the quality of recordings across the array at the time of implantation. The post-operative recovery regimen included administration of buprenorphine at 0.02 mg/kg and meloxicam at 0.2 mg/kg. Dexamethasone at 0.5 mg/kg and trimethoprim sulfadiazine at 15 mg/kg were also administered postoperatively for 5 days. All animals were allowed to recover for at least five days with the same regimen as described above before the start of experiments. Data collection and analysis were not performed blind to the conditions of the experiments.

### Photothrombotic and endothelin-1 induced focal stroke_exp_

We used either photothrombotic (PT) and Endothelin-1 (ET-1) induced stroke_exp_ models (see stroke_exp_ types for each group of animals in Table S1). For the PT stroke_exp_ model, rose bengal dye was injected into a femoral vein using and an intravenous catheter after craniotomy. Next, the surface of the brain was illuminated with white light (KL-1500 LCD, Schott) using a fiber optic cable for 20 min. We used a 4-mm aperture for stroke_exp_ induction (centered in the M1 area based on stereotactic coordinates; –1.5-2.5 mm anterior and 1-5 mm lateral from the bregma) and covered the remaining cortical area with a custom aluminum foil mask to prevent light penetration. After induction, a probe was implanted in the PLC immediately anterior to the lesion site (Gulati et al., 2015; Ramanathan et al., 2018). The craniotomy and implanted electrodes were covered with a layer of low toxicity silicone adhesive (WPI KWIK-SIL), then covered by dental cement. For the precise measure of baseline of LFP/spike activity before inducing stroke_exp_ (pre-stroke_exp_ baseline in Figure 7A), we also used ET-1 induced focal stroke_exp_ model (Robinson et al., 1990; Roome et al., 2014; Sharkey et al., 1993; Virley et al., 2004) in three rats (Figure 7). To induce ET-1 stroke_exp_, microarray attached with infusion cannula (guide: 26G; internal: 33G; P1 Technology) was implanted into the M1 area; single cannula in one rat and bilateral cannula with 2 mm spacing in the other two rats. The cannula was centered at 3 mm lateral and 0.5 mm anterior from the bregma to target M1 and microarray was positioned anterior to the cannula; the lesion site and recording site were targeted to correspond to the PT stroke_exp_ animals. The remaining operation procedures were the same as the PT stroke_exp_ animals. The ET-1 induced stroke_exp_ model allowed us to induce the focal stroke_exp_ following a baseline measurement; after measuring baseline spike/LFP activity for 2 days, ET-1 was injected with 1.5 ul through a single cannula in one rat, and a total 2.0 ul (two sites injections; 1.0 ul for each) through a bilateral cannula (100 nl per min) in the other two rats into deep cortical layers (1.4 mm from the surface of the brain). We then measured spike/LFP activity during the recovery period following the ET- 1 stroke_exp_ induction.

### Electrophysiology

We conducted AC-coupled recordings and recorded extracellular neural activity using 32-channel microwire electrode arrays (MEAs; 33-μm-length, 250-μm-spacing, 4-rows, standard polyimide-coated tungsten microwire arrays from Tucker-Davis Technologies (TDT) for ten rats; 25-μm- length, 200-μm-spacing, 6-rows, tungsten microwire arrays from Innovative Neurophysiology Inc. for five rats) and 128-channel custom electrode arrays (Egert D, 2018) (for one rat). All electrode arrays showed similar quality of LFP (e.g., LFP amplitude and noise level). The microwire arrays from Innovative Neurophysiology Inc. were customized so that a cannula is placed beside the recording sites of microwires. We recorded spike and LFP activity using a 128-channel RZ2 bioamp processor (TDT) and 128-channel neurodigitizer (digital PZ4 or PZ5).

Spike data was sampled at 24,414 Hz and LFP data at 1,018 Hz. ZIF-clip-based headstages (TDT) for 32-channel microwires and SPI-based headstates (Intan Technologies) for 128-channel microwire with a unity gain and high impedance (∼1 G) was used. Only clearly identifiable units with good waveforms and a high signal-to-noise ratio were used. The remaining neural data (e.g. filtered LFP) was recorded for offline analysis at 1,018 Hz. Behavior-related timestamps (that is, trial onset and trial completion) were sent to the RZ2 analog input channel using a digital board and then used to synchronize to neural data in the offline analyses. Electrophysiology was not monitored during the baseline sessions illustrated for the pre-stroke_exp_ motor performance in Figure 1 and in the animal group of Figure 4 (see Table S1 for the corresponding animals).

### Behavior

After recovery, animals were typically handled for several days before the start of experimental sessions, i.e., “motor training sessions.” Animals were acclimated to a custom plexiglass behavioral box during this period without motor training. The box was equipped with a door at one end. We examined two groups of stroke_exp_ animals; with motor training in Figures 1, 2, 3, 4, 5, and 6 and with spontaneous recovery in Figure 7. In the rats used for Figure 7 (testing the effect of reducing tonic GABA_A_), animals experienced spontaneous recovery without motor training. In the other rats of Figures 1, 2, 3, 4, 5, and 6, animals were trained to a plateau level of performance in a reach-to-grasp single-pellet task before neural probe implantation or PT stroke_exp_ induction (baseline; pre-stroke_exp_ training period in Figure 1A). We measured relative reach performance, i.e., pellet retrieval success rate in the reach-to-grasp task, using normalized metrics relative to the baseline, Figure 1B. However, the absolute reach performance was compared to sleep metrics for all other analyses. The reach-to-grasp task has been used as a sensitive measure of motor function; it requires reaching, grasping, and retrieving a single pellet located at a distance outside of the behavior box (Figure S1A) (Guo et al., 2021; Ramanathan et al., 2018; Whishaw et al., 1986; Wong et al., 2015). Probe implantation and stroke_exp_ induction were performed contralateral to the preferred hand. Animals were allowed to rest at least for 6 days before the start of motor training or recording sessions post-stroke_exp_. The stroke_exp_ animal group experienced motor training until the motor performance reached a plateau level of performance (motor performance in an individual animal is shown in Figure S1), which is called motor training post-stroke_exp_ during the recovery period in Figure 1A. The first reach training session for the “across-session” analyses was between the 7-14th day (9.1±2.6, mean±s.d.); this was because of the need to restart food scheduling. Each animal was monitored for 6-11 sessions/days until a stable plateau was reached (Figure S1). Given the variability of recovery times, and as described in the results, sessions per animal were divided into tertiles and termed ‘early’, ‘middle’ and ‘late.’ During behavioral assessments, we monitored the animals and ensured that their body weights did not drop below 90% of their initial weight. We monitored electrophysiology, i.e., LFP/spike only during the pre-training sleep, reach training, post-training sleep, and retrieval period of each session, not during 24 hours of a single day. This typically totaled a period of 4-5 hours a day in the behavior box. After completing motor tasks and sleep sessions in the behavioral box, animals were placed in the home cage without electrophysiology monitoring.

For the behavioral task, we used an automated reach-box, controlled by custom MATLAB scripts and an Arduino microcontroller. This setup for the reach-to-grasp task required minimal user intervention, as described previously (Wong et al., 2015). Each trial consisted of a pellet dispensed on the pellet tray, followed by a beep indicating that the trial was beginning; this was followed by the door opening. Animals then had to reach their arm out, grasp and retrieve the pellet. A real-time “pellet detector” using an infrared detector centered over the pellet was used to determine when the pellet was moved, which indicated that the trial was over and the door was closed. All trials were captured by video, which was synced with the electrophysiology data using the Arduino digital output. The video frame rate was 30 Hz for six animals and 75 Hz for two animals. The reach performance (the number of accurate pellet retrieval/the total number of trials x 100 %) was determined manually from a recorded video. The reach performance was used not only as a measure of motor function recovery across sessions (Figure 1B) but also as a measure of offline gains that may be the result of memory consolidation from sleep (Figure 3C). As a measure of offline gain, we computed the next-day reach performance (i.e., a change in reach performance between Day X reach training and the following Day X+1 reach training; Figure 3C) and the within-session changes (i.e., reach performance changes after post-training sleep compared to before post-training sleep, Figure 4). These measurements allowed us to quantify offline gains and motor recovery either across sessions and within a session (see also below).

### GABA_A_ α5-subtype receptor inverse agonist treatment

Treatment with the GABA_A_ α5-subtype receptor inverse agonist (L655,708; “drug” condition in Figure 7) was initiated 6 days after ET-1 stroke_exp_ induction; previous studies reported that L655,708 promoted functional recovery 7 days after stroke_exp_ (Clarkson et al., 2010; He et al., 2019). About 0.5∼1 hour before sleep onset, rats received intraperitoneal/IP injections of vehicle (i.e., saline as the “sham”) or L655,708 (Sigma-Aldrich; 5 mg/kg, dissolved in dimethylsulphoxide/DMSO and then diluted 1:1 in 0.9% saline as the “drug” condition). Over the course of 6 days, we used either sham or drug every other day (see experiment flow in Figure 7A). To counterbalance the starting condition between sham and drug conditions in six animals, the sham was conducted on days 7, 9, and 11 for ET-1 induced animals (n = 3), and the drug was conducted on days 7, 9, and 11 for PT induced animals (n = 3).

### Analyses across-session versus within-session

In the stroke_exp_ experiment in Figures 1, 2, 3, 4, 5, and 6, animals performed the reaching task during recovery (6-11 days of training). For long-term recovery in the stroke_exp_ animal group, we specifically analyzed the “pre-training sleep” and the “reach training” over a ∼3 weeks recovery period. In the other experiment, to test the effect of reducing tonic GABA_A_ in Figure 7, we focused on spontaneous changes in sleep microarchitectures without motor training over a ∼2 weeks recovery period. These analyses of the changes over a long-term recovery period are termed “across-session” analyses. In each animal, the 1-2 weeks recovery period was divided into sextiles and a representative session in each sextile was used. The motor performance and the sessions used are marked in Figures S1B. In five animals, all monitored sessions were used (i.e., two sessions per period and total of six sessions were monitored). In three animals, representative sessions (two sessions per period) were selected in order to balance the number of sessions per animal, e.g., typically every other session was selected. In Figures 3 and 4, we also compared sleep metrics within a single-day session/experiment (i.e., “within-session” analyses). Sleep metrics (e.g., SO-Nesting) were measured as a pair of pre-training and post-training sleep in a single-day session, then the change from the pre-training to the post-training sleep was computed (e.g., ΔSO-Nesting). These metrics (e.g., Day X) were then compared with the offline gains in motor performance during the reach training of the following day (e.g., Day X+1) in Figure 3C. In Figure 4, the within-session offline gains in motor perfromance (i.e., changes from reach training to reach retrieval within a single session) were examined.

### Identification of NREM sleep waves

LFP activity was recorded using 32/128-channel microwire electrode arrays (see above). The LFP was analyzed after removing obvious artifacts (> 10 s.d.) and excluding bad channels. Identification of NREM epochs was performed by classification based on power spectral density of the LFP. LFP trace was segmented into non-overlapping 6-sec epochs. In each epoch, the power spectral density was computed and averaged over the slow-wave frequency band including SO and δ-waves (0.1-4 Hz, also called Delta band) and Gamma frequency bands (30-60 Hz). Then a k-means classifier was used to classify epochs into two clusters, NREM sleep and REM/awake; REM sleep and awake were not classified and NREM sleep was focused on in this study. Sleep epochs less than 30 sec were excluded from NREM sleep epochs. The identified NREM sleep durations were not different between the early period and late period of motor recovery (Figure S2B). The identified NREM sleep epochs were verified by visual assessment of the LFP activity. During the NREM period with high Delta power (0.1-4 Hz), strong down- and up-states dominates. Thus, we assessed if our detected NREM sleep epochs contain a high-amplitude and slow LFP fluctuation distinguished from a low-amplitude and high-frequency LFP during the awake period. Moreover, we assessed if there were substantial missing detections of NREM sleep epoch; assessment if a high amplitude LFP epoch was not included in the detected NREM sleep epochs excessively. These power-based sleep detections showed a close match to the video-based detections (Kim et al., 2019; Pack et al., 2007) (Figure S2A); the number of pixels that changed intensity frame to frame in each pair of consecutive frames was computed from a recorded video (1 Hz frame rate using Microsoft LifeCam Cinema Webcam) during the sleep block; these values were then integrated over an epoch of 40 sec. If that integrated value was higher than a threshold, that epoch was identified as sleep; the threshold was chosen by comparing detection results and visual assessment of the recorded video.

In offline analysis, SO, δ-waves and spindles were detected using the algorithm used in previous studies (Gulati et al., 2017; Kim et al., 2019; Lemke et al., 2021; Silversmith et al., 2020). The LFP average across all recording channels excluding bad channels was filtered in the SO/δ band (0.1-4 Hz) through two independent filterings; the high pass Butterworth filter (2nd order, zero phase-shifted, with a cutoff at 0.1 Hz) was applied and then followed by the low pass Butterworth filter (5th order, zero phase-shifted, with a cutoff at 4 Hz). The individual order of the high-pass and low-pass filter was estimated through a conventional minimum-order design; it requred to meet maximum passband ripple of 3 dB and minimum stopband (presumed 0.02 Hz for high-pass filter and 6 Hz for low-pass filter) attenuation of 15 dB. Next, all positive-to-negative zero crossings during NREM sleep were identified, along with the previous peaks, the following troughs, and the surrounding negative-to-positive zero crossings. Then the positive threshold (the top 15 percentile of the peaks) and the negative threshold (the bottom 40 percentile of the troughs) were respectively defined for the down-states and up-states. Each identified wave was considered a SO if the trough was lower than the negative threshold (i.e., up-state), the peak preceding that up-state was higher than the positive threshold (i.e., down-state), and the duration between the peak and the trough was between 150 ms and 500 ms (Figures 2A and 5A). On the other hand, a slow wave was considered a δ-wave if the trough was lower than the negative threshold (i.e., up-states) and that the up-state was preceded by a maximum voltage that was lower than the positive threshold within 500 ms. In this study, we separated δ-waves into two classes depending on their temporal interaction with a preceding SO, i.e., ΔT_SO-δ_ (Figures 5B and C). ΔT_SO-δ_ > 100 sec (1∼2%) were manually removed to exclude the cases driven by the possible error in NREM sleep detections; this error was evaluated by visual inspection of slow-wave activity. From the distribution of ΔT_SO-δ_ in healthy brains (n = 9 sessions, 5 rats), 50^th^ percentile of ΔT_SO-δ_ (3.7 sec) was set as the cutoff separating δ-waves into “SO-coupled δ/δ_SO_” and “isolated δ/δ_I_” waves; δ- waves within the close 50% from a preceding SO (ΔT_SO-δ_ ≤ 3.7 sec) were labeled “δ_SO_,” whereas δ-waves of the distal 50% from a preceding SO (ΔT_SO-δ_ > 3.7 sec) were labeled “δ_I_.” The cutoff of ΔT_SO- δ_ determined by the healthy conditions was also used for stroke_exp_ conditions. The distribution of ΔT_SO-δ_ was more variable across the stroke_exp_ animals; it appears to be due to the variability of lesion size. Specifically in the comparison with the same animal (Figure S6), the distribution of ΔT_SO-δ_ after stroke_exp_ appeared to be abnormal compared to the healthy condition without stroke_exp_. Therefore, the cutoff of ΔT_SO-δ_ was determined by using the healthy conditions.

For spindles detection, the LFP was first z-scored in each channel and averaged across all good channels. The LFP average was filtered in spindle band (10-15 Hz) through two independent zero phase-shifted filterings (Figures 2A and 6A); the high pass Butterworth filter (6th order, zero phase-shifted, with a cutoff at 10 Hz) was applied and then followed by the low pass Butterworth filter (8th order, zero phase-shifted, with a cutoff at 15 Hz). The individual order of the high-pass and low-pass filter was estimated through a conventional minimum-order design; it requred to meet maximum passband ripple of 3 dB, and minimum stopband (presumed 7 Hz for high-pass filter and 19 Hz for low-pass filter) attenuation of 15 dB. We computed a smoothed envelope of this signal, the magnitude of the Hilbert transforms with convolving by a Gaussian window (200 ms). Next, we determined two thresholds for spindle detection based on the mean (μ) and standard deviation (σ) of the spindle band LFP during NREM sleep; the upper and lower thresholds were set μ + 2.5 × σ and μ + 1.5 × σ, respectively. Epochs in which the spindle power exceeded the upper threshold for at least one sample and the spindle power exceeded the lower threshold for at least 500 ms were considered spindles. Each epoch where the spindle power exceeded the lower threshold was considered the start and stop of the spindle; the duration of each spindle was based on these values as well.

### Spindle nesting analyses

We also analyzed the temporal coupling of spindles relative to SO or δ-waves. For the nesting of spindles to SO (SO-Nesting; Figure 2B), each spindle was linked to the closest SO. The time difference between the peak of the spindle and the up-state of the linked SO was measured for each detected spindle (ΔT_SO-Spindle_). If ΔT_SO-Spindle_ was between –0.5 sec and 1.0 sec (i.e., nesting time window), that spindle event was considered a SO-nested spindle. In this study, we also focused on the temporal interaction of spindles to the δ_I_-waves. Thus, the nesting of spindles to δ_I_ (δ_I_-Nesting; Figure 5D) was identified in a manner analogous to the “SO-Nesting” value, i.e., time differences between the spindle peak time and the time of the δ_I_ up-state. To quantitatively assess the changes in the temporal coupling of spindles to SO, we specifically measured as the following; time lag of spindle from the closest SO (ΔT_SO-Spindle_) was measured for each spindle event and the rate of spindles of which ΔT_SO-Spindle_ was within the nesting time window was measured; i.e., the number of SO-nested spindles / the total number of spindles × 100 % (Figures 2E and F). We also measured δ_I_-Nesting the same way as the “SO-Nesting” but using time lags between the peaks of spindles and the up-states of the linked δ_I_; i.e., the number of δ_I_-nested spindles / the total number of spindles × 100 % (Figures 5E, and F).

### Spike activity during sleep waves

We initially used an online sorting program (SpikePac, TDT). We then conducted offline sorting using Plexon Inc’s Offline Sorter. Only clearly identifiable units with good waveforms and a high signal-to-noise ratio were used. To assess spike activity modulation during sleep oscillations, we analyzed peri-event time histogram (PETH) and unit modulation-depth (MD). After spikes were time-locked to event reference times (e.g., SO up-states or SO-nested spindles), the PETH (bin length 20 ms) was estimated. Then, the unit MD was calculated by comparing the difference between maximum and minimum of the PETH around events time (within –0.5 to 0.3 sec from SO up-states or –0.25 to 0.15 sec from spindles peaks) over the baseline firing activity (averaged activity within –2 to –0.5 sec from the events); i.e., (maximum-minimum)/baseline firing rate. In other words, the MD is a measure of the modulation of firing rate relative to the pre-event start baseline rate. This was compared for the early period and the late period of recovery in Figure 2G and for the sham and drug conditions in Figures 7C and D. No significant change in population spike rates was found from the early to the late period (Figure S4A).

### Quantification and statistical analyses

Figures show mean ± s.e.m.; if this was not the case, we specifically indicated it. Parametric statistics were generally used in this study (linear mixed-effect model, *t*-tests, linear regression, Pearson’s correlation otherwise stated); they were implemented within MATLAB. A “hierarchical nested statistics approach” of linear mixed-effects models (using the MATLAB function “fitlme”) was used for the comparison of task performance, temporal coupling of spindles, unit modulation-depth, and linear relationship (Aarts et al., 2014; Guo et al., 2021; Khanna et al., 2021; Kim et al., 2019). This was done to account for the repeated measures per animals; thus, this statistical approach ensured that the group level statistc accounted for sessions per animal and did not treat them as statistically independent samples. We fit random effects (e.g., rats) specified as an intercept for each group and reported fixed effects representing population parameters to compare (e.g., early vs late period). Adding random effects to a model recognizes correlations within sample subgroups (e.g., rat) and extends the reliability of inferences beyond the variability across multiple rats. The fixed effects were tested for P values of the linear regression slope coefficients associated with two comparing conditions. In this way, the mixed-effects model accounts for the fact that units, sessions, events, or experimental conditions from the same animal are more correlated than those from different animals and is more stringent than computing statistical significance over all units, sessions, events, and conditions. The mixed-effects model was used to compare stroke_exp_ versus healthy conditions, early period versus late period, sham versus drug, and sleep versus no-sleep. The used random effects and fixed effects parameters are following; Figures 2F, 2G, 3B, and 5F, and Figures S2, S3, S4, and S5, random: rat, fixed: recovery period; Figure 4A, random: rat, fixed: sleep type; Figures 7B and C, random: rat, fixed: injection type. In these figures, the mean in each experiment session was used as the response parameter and two categories of the comparing conditions were used as the predictor parameter.

The mixed-effects model was also used to evaluate linear relationships or correlations between the changes in SO-Nesting and the next-day task performance in Figure 3C, between the post-training sleep duration and the within-session changes of task performance in Figure 4B, between the mean rate of δ_I_ preceding SO and the SO-Nesting in Figure 6B, between the mean rate of δ_I_ preceding SO and the next-day task performance in Figure 6C, and between the changes of SO-Nesting and the changes in spindles-MD in Figure 7D. We also used traditional linear regression or correlation to evaluate the relationship between the recovery period and the sleep metrics (e.g. SO-Nesting and δ_I_-Nesting) in Figures 2E and 5E, and Figure S3 left. For the comparison between distributions, we used Kolmogorov-Smirnov’s test in Figure 5C and Figure S6 to test the samples were drawn from the same distribution and Bartlett’s test in Figure 2D to test equal variances (sharpness of distribution) between early period and late period.

## Acknowledgements

Research was supported by awards from the Department of Veterans Affairs, Veterans Health Administration (VA Merit: 1I01RX001640 to K.G.; VA CDA to DR, 7IK2BX003308); the Weill Neurohub, the National Institute of Neurological Disorders and Stroke (5K02NS093014 to K.G.; K99NS119737 to J.K.); American Heart Association Postdoctoral Fellowship (831442 to J.K.); and the Basic Science Research Program through the National Research Foundation of Korea (2018R1A6A3A03013031 to J.K.). Karunesh Ganguly, M.D., Ph.D., and Dhakshin Ramanathan, MD., PhD hold a Career Award for Medical Scientists from the Burroughs Wellcome Fund.

## Author Contributions

JK, LG, DSR, SW and KG conceived and designed the experiments. JK, LG, AH, SL, and SW conducted the experiments. JK analyzed all data, LG analyzed the data for Figure 1B, and AH analyzed the data for Figure 4. JK and KG wrote and edited the manuscript.

