## Supplementary Information for "Recovery of consolidation after sleep following experimental stroke – interaction of slow waves, spindles and GABA"

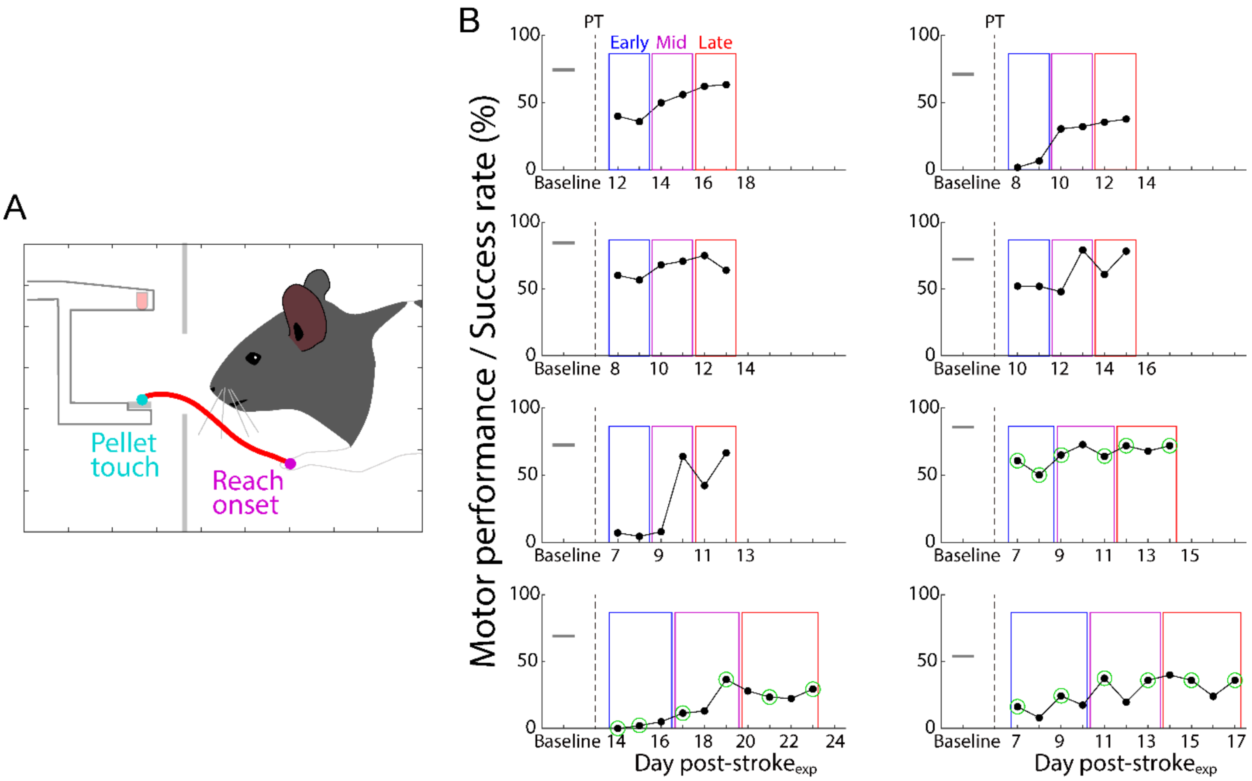


**Figure S1. Reach-to-grasp task and success rates over all recovery periods.**

**(A)**, Cartoon example of reach-to-grasp motor task with lateral view. Mageta and cyan dots represents reach onset and pellet touch position, respectively. Red trace represents an example of mean trajectory within a single session.

**(B)**, Pellet retrieval success rates in an individual animal (n=8 rats). Three colored boxes repsents the presumed tertile recovery periods after induction of PT stroke$_{exp}$. In three rats in which motor recovery was monitored more than 6 days, we used two representative sessions per period (green circles).

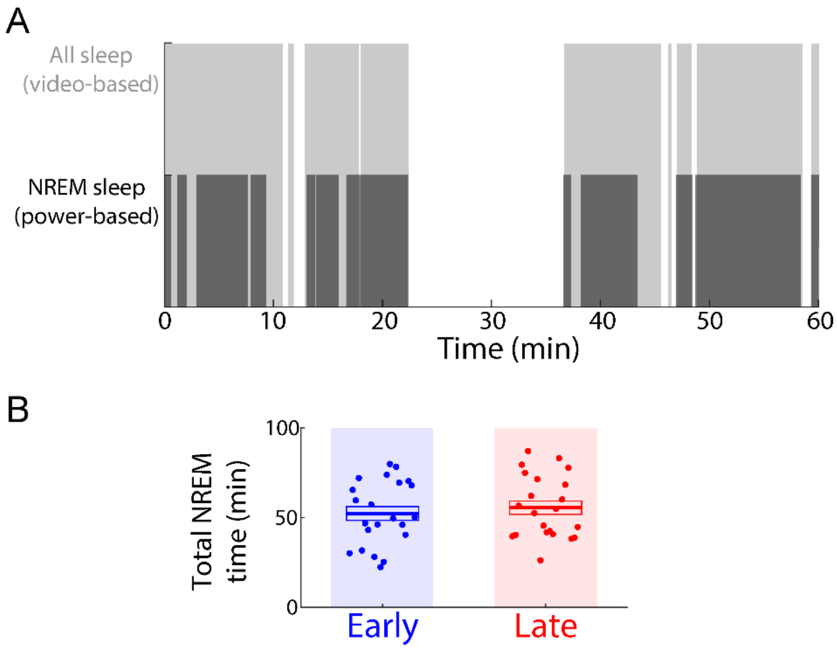


**Figure S2. Sleep detection and NREM sleep duration.**
**(A)**, Examples of detections of all sleep using a degree of movement (gray; see Methods) and NREM sleep using power spectral density in 0.1-4 Hz and gamma (30-60 Hz) bands (black).
**(B)**, Mean NREM sleep time during early and late period after stroke. NREM sleep time across all sessions: 52.2 ± 3.9 min during early period (n = 28 sessions in 14 rats), 55.5 ± 3.7 min during late period (n = 28 sessions in 14 rats). No significant difference; early vs. late: mixed-effects model, $t_{54} = 0.93$, $P = 0.36$. Mean in solid line ± s.e.m. in box.

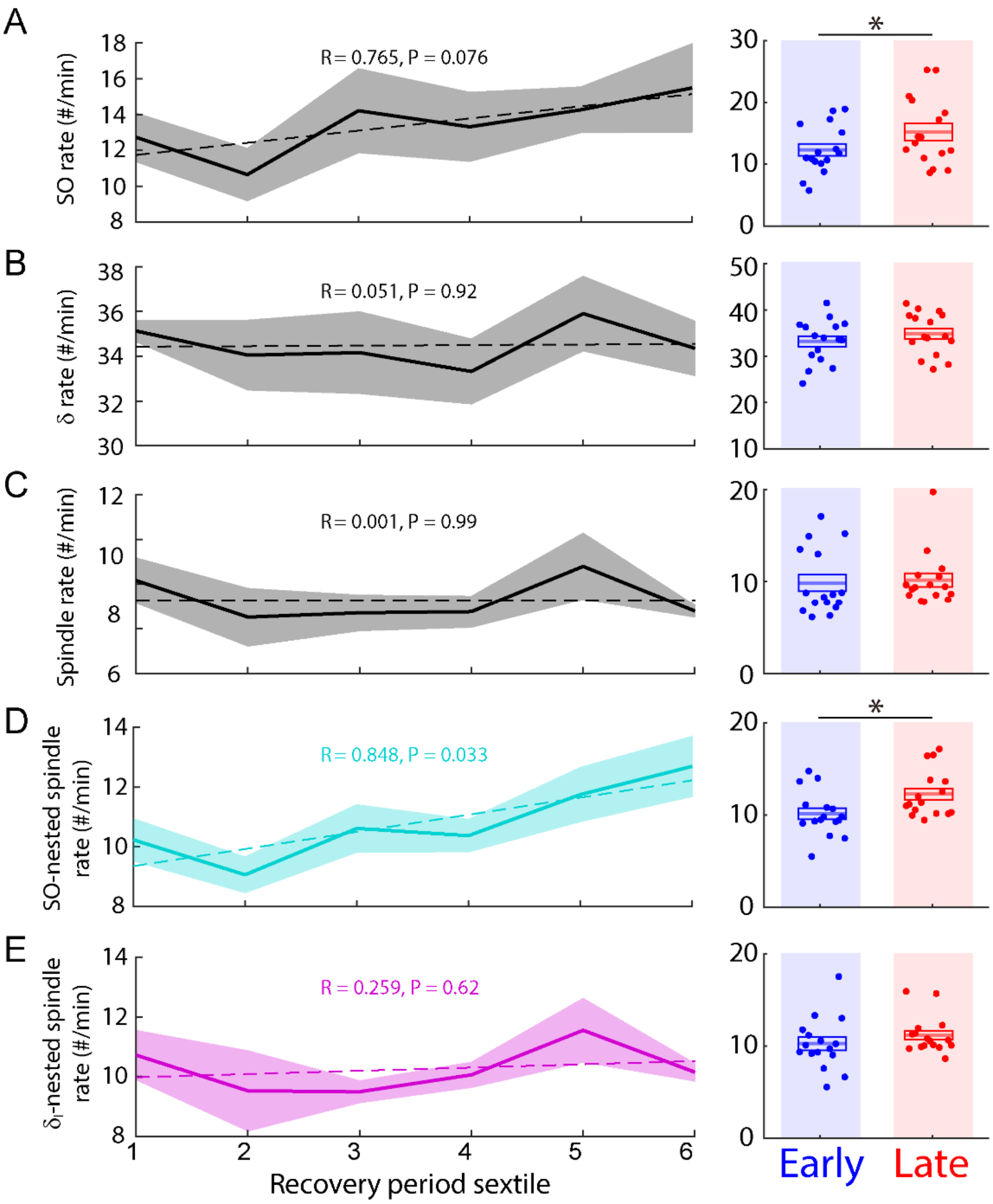


**Figure S3. Sleep waves rates over the recovery period.**
Left, average time courses in the rates of SO (**A**), δ-waves (**B**), spindles (**C**), SO-nested spindles (**D**), and $\delta_l$-nested spindles (**E**) over the recovery period sextile (n = 11 rats). Dashed line represents linear fitting (linear regression; SO: R = 0.765, P = 0.076; δ: R = 0.051, P = 0.92; spindles: R = 0.001, P = 0.99; SO-nested spindles: R = 0.848, P = 0.033; $\delta_l$-nested spindles: R = 0.259, P = 0.62). Right, comparisons between early period and late period in each corresponding metric (early vs. late, mixed-effect model; SO: $t_{30}$ = 2.21, *P = 0.035; δ-waves: $t_{30}$ = 1.65, P = 0.11; spindles: $t_{30}$ = 0.28, P = 0.77; SO-nested spindle: $t_{30}$ = 2.52, *P = 0.017; $\delta_l$-nested spindles: $t_{30}$ = 1.18, P = 0.25). Mean in solid line ± s.e.m. in box.

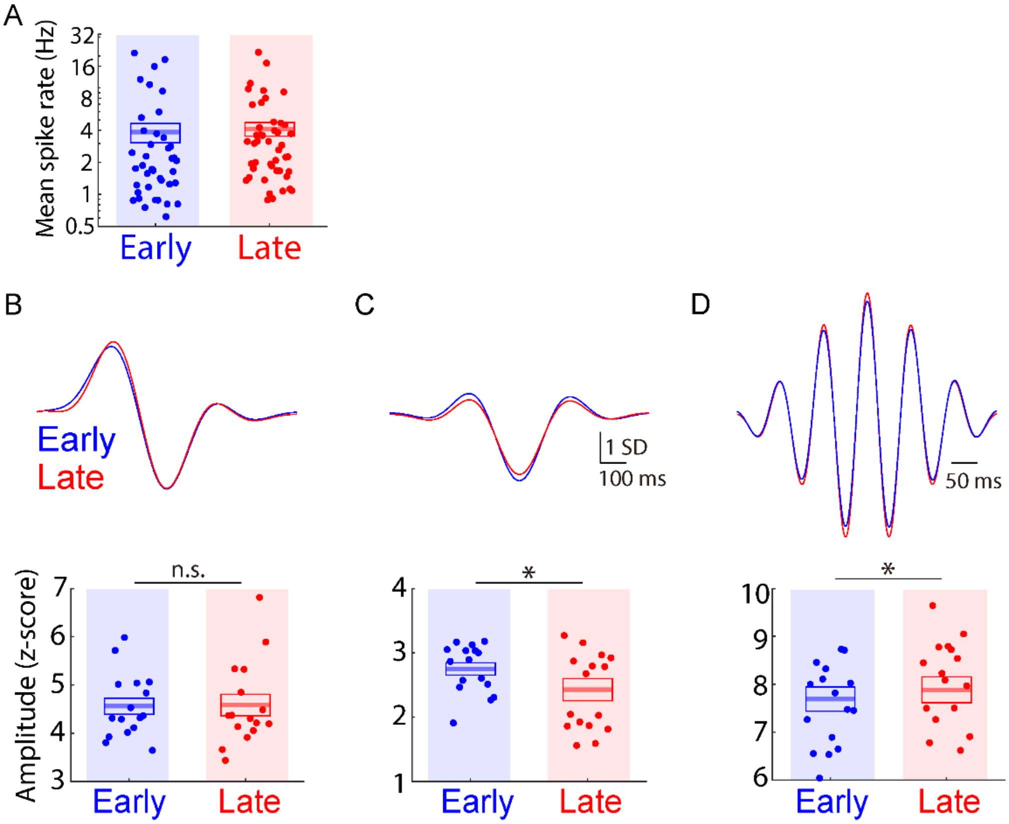


**Figure S4. Spike activity and Sleep waves amplitude.**
**(A)**, Mean spike rates of units during NREM sleep for early period and late period after $stroke_{exp}$. There was a similar activity for both periods (mixed-effects model, $t_{97} = -0.29$, P = 0.77).
**(B-D)**, Top, averages of event-triggered LFPs of SO (**B**), $\delta_l$ (**C**), and SO-nested spindles (**D**) for early period (blue; n = 16 sessions, 8 rats) and late period after $stroke_{exp}$ (red; n = 16 sessions, 8 rats). LFP was normalized; subtracting by the mean and dividing by the standard deviation in each sleep session. Bottom, mean peak-to-trough amplitude corresponding to top. There were significant changes for $\delta_l$ and SO-nested spindles from early period to late period (SO: mixed-effects model, $t_{30} = 0.13$, P = 0.90; $\delta_l$: mixed-effects model, $t_{30} = -2.60$, *P = 0.014; SO-nested spindles: mixed-effects model, $t_{30} = 2.26$, *P = 0.031). Mean in solid line ± s.e.m. in box.

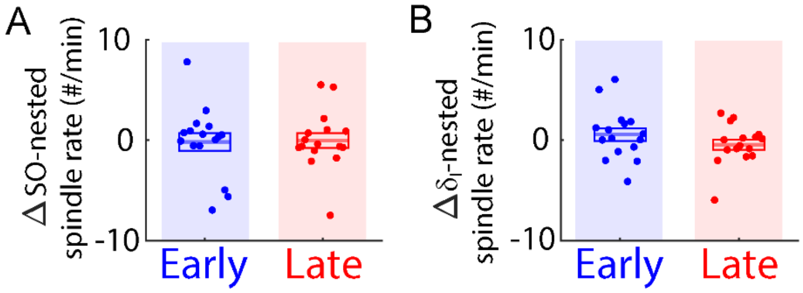


**Figure S5. Changes in spindles' rates within a single session**

Changes of rates in the SO-nested spindles (**A**) and the $\delta_l$-nested spindles (**B**) within a single session (i.e., from pre-training to post-training sleep) in each column of the two recovery periods. ΔSO-nested spindles (early, n = 16 sessions in 8 rats, –0.60 ± 0.45% vs. late, n = 16 sessions in 8 rats, 0.33 ± 0.61%, mixed-effects model, $t_{30}$ = 1.27, P = 0.21). Δ$\delta_l$-nested spindles (early, n = 16 sessions in 8 rats, 0.56 ± 0.63% vs. late, n = 16 sessions in 8 rats, –0.43 ± 0.50%, mixed-effects model, $t_{30}$ = –1.34, P = 0.19). Mean in solid line ± s.e.m. in box.

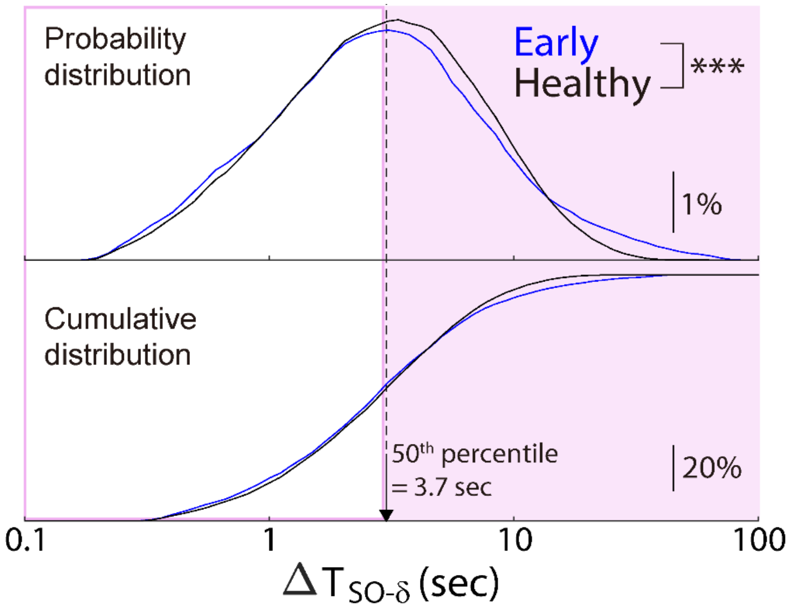


**Figure S6. Redistribution of $\delta_I$-waves relative to SO after $stroke_{exp}$ in the same animal.** Example distributions of $\Delta T_{SO\text{-}\delta}$ are shown for sleep sessions in the early recovery (blue; n = 1 rats) and sleep sessions in the intact brain before inducing ET-1 $stroke_{exp}$ in the same rat. (black; n = 1 rat; Kolmogorov-Smirnov test, KS-statistic = 0.18, ***$P < 10^{-9}$).

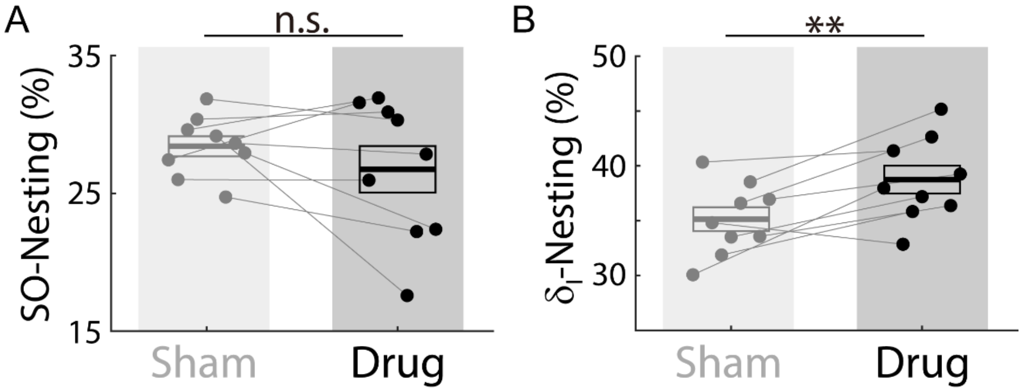


**Figure S7. Blocking $GABA_A$ α5-subtype receptor in healthy animals.**
Comparison of the drug (L655,708) effect on SO-Nesting **(A)** and $\delta_l$-Nesting **(B)** in healthy animals. SO-Nesting was not changed while $\delta_l$-Nesting was stronger, with the drug compared to the sham (SO-Nesting: sham, n = 9 sessions, 28.4 ± 0.73% vs. drug, n = 9 sessions, 26.8 ± 1.7%, mixed-effects model, $t_{16} = -1.14$, $P = 0.27$; $\delta_l$-Nesting: sham, 35.2 ± 1.1% vs. drug, 38.7 ± 1.3%, mixed-effects model, $t_{16} = 3.59$, $**P < 10^{-2}$).

| | Rat ID | Figures 1, 2, 3, 5, and 6 | | | | Figure 1 | Figure 7 | | | Figure 4 | |
|---|---|---|---|---|---|---|---|---|---|---|---|
| | | stroke$_{exp}$ + motor training | | | | Healthy | stroke$_{exp}$ + spontaneous recovery | | | stroke$_{exp}$ + motor training | |
| | | Baseline | Early | Middle | Late | | Baseline | Sham | Drug | Sleep | No-sleep |
| Contribution (number of sessions) | 1 | $2^{\#}$ | 2 | 2 | 2 | - | - | - | - | - | - |
| | 2 | $2^{\#}$ | 2 | 2 | 2 | - | - | - | - | - | - |
| | 3 | $2^{\#}$ | 2 | 2 | 2 | - | - | - | - | - | - |
| | 4 | $2^{\#}$ | 2 | 2 | 2 | - | - | - | - | - | - |
| | 5 | $2^{\#}$ | 2 | 2 | 2 | - | - | - | - | - | - |
| | 6 | $2^{\#}$ | 2 | 2 | 2 | - | - | - | - | - | - |
| | 7 | $2^{\#}$ | 2 | 2 | 2 | - | - | - | - | 5 | - |
| | 8 | $2^{\#}$ | 2 | 2 | 2 | - | - | - | - | 5 | - |
| | 9 | - | - | - | - | 2* | 2* | 3 | 3 | - | - |
| | 10 | - | - | - | - | 2* | 2* | 3 | 3 | - | - |
| | 11 | - | - | - | - | 2* | 2* | 3 | 3 | - | - |
| | 12 | - | - | - | - | - | - | 3 | 3 | - | - |
| | 13 | - | - | - | - | - | - | 3 | 3 | - | - |
| | 14 | - | - | - | - | - | - | 2 | 2 | - | - |
| | 15 | - | - | - | - | 2 | - | - | - | - | - |
| | 16 | - | - | - | - | 1 | - | - | - | - | - |
| | 17 | - | - | - | - | - | - | - | - | $6^{\#}$ | - |
| | 18 | - | - | - | - | - | - | - | - | $5^{\#}$ | - |
| | 19 | - | - | - | - | - | - | - | - | $5^{\#}$ | - |
| | 20 | - | - | - | - | - | - | - | - | $4^{\#}$ | - |
| | 21 | - | - | - | - | - | - | - | - | - | $6^{\#}$ |
| | 22 | - | - | - | - | - | - | - | - | - | $4^{\#}$ |
| | 23 | - | - | - | - | - | - | - | - | - | $4^{\#}$ |
| | 24 | - | - | - | - | - | - | - | - | - | $2^{\#}$ |
| | 25 | - | - | - | - | - | - | - | - | - | $3^{\#}$ |
| | 26 | - | - | - | - | - | - | - | - | - | $4^{\#}$ |
| | 27 | - | - | - | - | - | - | - | - | - | $3^{\#}$ |
| | 28 | - | - | - | - | - | - | - | - | - | $2^{\#}$ |
| | Total | 16 | 16 | 16 | 16 | 9 | 6 | 17 | 17 | 29 | 28 |

**Table S1. The number of sessions contributed by each rat to the respective experiment types.**

All numbers indicate independent contributions to each condition except the numbers marked by star; * indicate the same session that is counted for two conditions. # indicates the sessions in which electrophysiology was not monitored. Rat 9-16 was monitored only for sleep without behavior tasks. Stroke$_{exp}$ models: photothrombotic/PT for Rat 1-8, 12-14, and 17-28, and ET-1 induced for Rat 9-11.
